# Detection of a Mitochondrial Stress Phenotype using the Cell Painting Assay

**DOI:** 10.1101/2023.11.08.565491

**Authors:** Soheila Rezaei Adariani, Daya Agne, Sandra Koska, Annina Burhop, Jens Warmers, Petra Janning, Malte Metz, Axel Pahl, Sonja Sievers, Herbert Waldmann, Slava Ziegler

## Abstract

Mitochondria are cellular powerhouses and crucial for cell function. However, these organelles are vulnerable to internal and external perturbagens that may impair mitochondrial function and eventually lead to cell death. In particular, small molecules may impact mitochondrial function and cardio- or hepatotoxicity caused by numerous drugs links mitochondrial toxicity to these adverse effects. Therefore, the influence of small molecules on mitochondrial homeostasis is at best assessed early on in the characterization of biologically active small molecules and drug discovery. We demonstrate that unbiased morphological profiling by means of the Cell Painting assay (CPA) can detect mitochondrial stress coupled to the induction of integrated stress response. This activity is common for compounds addressing different targets, is not shared by direct inhibitors of the electron transport chain and enables prediction of mitochondrial stress induction for small molecules that are profiled using CPA.

## Introduction

Mitochondria are multifunctional signaling organelles that are essential for cellular homeostasis. They are involved in numerous vital processes beyond oxidative phosphorylation (OXPHOS) and ATP production, like lipid oxidation, one carbon metabolism and pyrimidine biosynthesis, ion uptake, synthesis of Fe/S clusters, anaplerosis, catabolism of various substrates, redox homeostasis, cell signaling etc. (Monzel et al., 2023). Mitochondria are linked to aging and diseases such as diabetes type 2, cardiovascular and Alzheimer’s diseases (San-Millán, 2023). Small molecules can selectively modulate mitochondrial targets and processes (Jalal et al., 2015). However, drug-induced mitochondrial toxicity has been linked to adverse effects such as cardio-, renal and hepatotoxicity as the heart and kidney are highly aerobic organs and the liver faces high drug concentrations due to drug accumulation (Dykens and Will, 2007). This adverse effect may result from inhibition of mitochondrial DNA (mtDNA) synthesis and electron transport chain (ETC), uncoupling of the proton gradient, induction of oxidative stress or mitochondrial permeability (Dykens and Will, 2007). Different approaches have been employed to assess impairment of mitochondrial function such as cell growth studies in presence of glucose (Glu) or galactose (Gal) (i.e., Glu/Gal assays (Marroquin et al., 2007)), metabolic flux measurements or detection of the mitochondrial membrane potential. The characterization of new small-molecule tools and drug candidates would tremendously benefit from detection of mitochondrial impairment early in the compound discovery process.

Here we report on the use of unbiased morphological profiling by means of the Cell Painting assay (CPA) (Gustafsdottir et al., 2013) for the detection of mitochondrial stress response upon compound treatment in U-2OS cells. In CPA, cells are stained with six different dyes for detection of cell organelles and components (DNA, RNA, mitochondria, Golgi, plasma membrane, endoplasmic reticulum, actin cytoskeleton) (Bray et al., 2016; Gustafsdottir et al., 2013). The obtained profiles are then compared with the profiles of reference compounds, i.e., compounds with known targets or modes of action (MoAs) and profile similarity can be used for the generation of target or MoA hypotheses (Akbarzadeh et al., 2022; Schneidewind et al., 2020; Schneidewind et al., 2021; Scholermann et al., 2022; Ziegler et al., 2021). We identified several small molecules with different targets or mechanism of action sharing similar morphological profiles, which is induced by impairment of mitochondrial function. This phenotype is detected for some but not all tested iron chelators, is linked to increased levels of mitochondrial superoxide, suppression of mitochondrial respiration after long-term treatment and activation of cyclic AMP-dependent transcription factor 4 (ATF4) and therefore, integrated stress response (ISR). Based on a set of similar CPA profiles related to mitochondrial stress, a consensus subprofile for a ‘MitoStress’ compound cluster was extracted according to a recently described approach (Pahl et al., 2023). For all CPA-active reference compounds tested by us, biosimilarity to this cluster can easily be assessed via the web app tool https://cpcse.pythonanywhere.com/, which will support interpretation of results after small-molecule treatment of cells regarding mitochondrial function and may guide the use and prioritization of bioactive small molecules for cell-based studies.

## Results

### Analysis of the CPA profiles of ciclopirox

We have screened 4,251 reference compounds and more than 10,000 in-house compounds using CPA (Akbarzadeh et al., 2022; Pahl et al., 2023; Schneidewind et al., 2020; Schneidewind et al., 2021; Scholermann et al., 2022). U-2OS cells were exposed to the compounds for 20 h, followed by staining of cell compartments and components including the plasma membrane, actin, DNA, RNA, the Golgi, the endoplasmic reticulum and mitochondria (Bray et al., 2016; Gustafsdottir et al., 2013). High-content imaging and analysis resulted in profiles composed of 579 features, which are Z scores representing the differences to the DMSO control (Christoforow et al., 2019). To describe activity in CPA, we use an induction value (in percent) which is the number of features that are significantly altered compared to the DMSO control and compounds are considered active for induction ≥ 5 %. Profile similarity (biosimilarity, BioSim, in %) is calculated based on Pearson’s correlation and profiles are similar if BioSim ≥ 75 %. Compounds with biosimilar CPA profiles are expected to share the same target or mode of action (MoA) and can be employed for the generation of target or MoA hypotheses. Our previous analysis of CPA profiles led to the definition of thus far twelve bioactivity clusters (Pahl et al., 2023) that are based on compound profile similarity, irrespective of different target annotation (Akbarzadeh et al., 2022; Pahl et al., 2023; Schneidewind et al., 2020; Schneidewind et al., 2021; Scholermann et al., 2022). To simplify the target or MoA prediction using CPA, we recently introduced the concept of subprofile analysis (Pahl et al., 2023). For this, features altered in the same direction are extracted from the full profiles recorded for biosimilar compounds that define each cluster. Using the reduced set of features for these compounds, a median profile is generated, termed cluster subprofile, which can be used to calculate cluster biosimilarity (Pahl et al., 2023).

As recently reported, the CPA profile for the metal ion chelator ciclopirox shares biosimilarity to the profiles of compounds that impair DNA synthesis at a concentration of 10 µM (Schneidewind et al., 2020). However, the profiles of ciclopirox at 30 and 50 µM are not biosimilar to the profile at 10 µM (Figure 1A). This is in line with a dose-dependent decrease in similarity to the DNA synthesis cluster (Figure 1B). The ciclopirox profiles at 30 and 50 µM were not similar to subprofiles of the remaining eleven clusters pointing towards a thus far unexplored bioactivity in CPA. We compared the profiles of ciclopirox and other iron-chelating agents like deferoxamine (DFO) and phenanthroline (Schneidewind et al., 2020). The profiles of DFO (3-30 µM) and phenanthroline (10 µM) define the DNA synthesis cluster (Pahl et al., 2023). The profiles recorded for DFO are biosimilar to each other at all tested concentrations and similar observations were made for phenanthroline (Figure S1A and S1B). In line with this, the profiles of DFO and phenanthroline were biosimilar to the DNA synthesis cluster at all tested concentrations (up to 50 µM) (Figure S1C and Figure 1D). No biosimilarity was observed for the profiles of DFO and 50 µM ciclopirox, whereas profile biosimilarity of 75 % was detected for 50 µM phenanthroline and 50 µM ciclopirox (Figure 1D). Thus, iron chelators share similar CPA profiles, however, morphological differences are detected at higher concentrations hinting to dose-dependent phenotype shift for some iron chelators.

**Figure 1.**
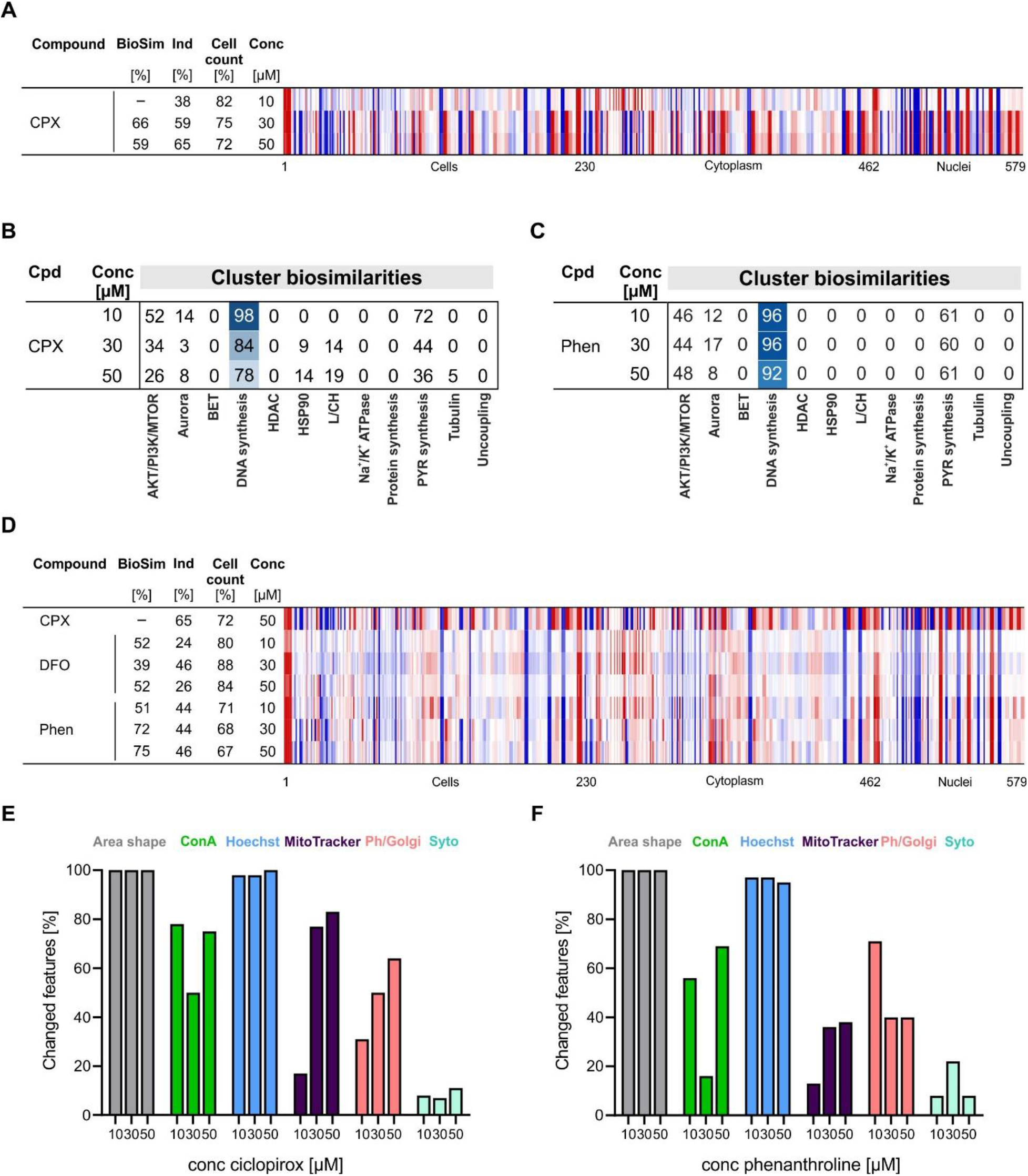
CPA profiles of ciclopirox. (A) Comparison of the profiles of ciclopirox (CPX) at different concentrations. The top line profile is set as a reference profile (100 % biological similarity, BioSim) to which the following profiles are compared. Blue color: decreased feature; red color: increased feature. (B) Cluster biosimilarity heatmap for ciclopirox. Percent values are given. (C) Cluster biosimilarity heatmap for the profiles of phenanthroline. Percent values are given. (D) Comparison of the profiles of DFO, phenanthroline and 50 µM ciclopirox. The top line profile is set as a reference profile (100 % biological similarity, BioSim) to which the following profiles are compared. Blue color: decreased feature; red color: increased feature. (E and F) Dose-dependent change in dye-related CPA features recorded for ciclopirox (E) or phenanthroline (F). Cpd: compound; BioSim: biosimilarity; Ind: induction; Conc: concentration. See also Figure S1.

To gain more information on the type of morphological changes, we compared the altered features in the profiles of ciclopirox at 10, 30 and 50 µM that are related to the individual CPA stains. All Hoechst-related features were changed at the three concentrations, which is related to the impairment of DNA synthesis and the cell cycle (Figure 1E) (Schneidewind et al., 2020). However, the number of changed MitoTracker features increased dose-dependently (Figure 1E) pointing towards impairment of mitochondria. To a smaller extent, a similar trend was observed for phalloidin/WGA-related features (Figure 1F). In contrast, treatment with DFO and phenanthroline did not change the MitoTracker-related features in a similar way (Figure 1F and Figure S1D). The altered features with the highest Z-scores for 30 µM ciclopirox were only MitoTracker-related features (see Figure S1E and Table S1). Hence, high concentrations of ciclopirox alter mitochondrial morphology.

### Similarity to the profiles of reference compounds

Exploring the profiles of reference compounds that are biosimilar to the ciclopirox profile at 30 µM revealed several biosimilar reference compounds like the Hypoxia Inducible Factor (HIF) pathway activator ML228, the gp130 (IL-6β) inhibitor SC144 (Lu et al., 2020), the dual JMJD3/KDM6B and UTX/KDM6A inhibitor GSK-J4, the natural products sanguinarine and chelerythrine, the ALK5 inhibitor **1** (Gellibert et al., 2009) and the anthelmintic agent pyrvinium pamoate (Table S2, Figure 2A). These compounds differ in their target annotation (see Table S2). Some of them modulate their targets by chelating metal ions, e.g., ML228 and GSK-J4 (Kruidenier et al., 2012; Theriault et al., 2012) and most likely SC144, whose cytotoxicity could be rescued by supplementation of iron, copper or zinc ions (Lu et al., 2020). Furthermore, diverse activities have been reported for sanguinarine, chelerythrine and pyrvinium pamoate (Chen et al., 2022; Croaker et al., 2016; Ishii et al., 2012). Therefore, the profiles of these compounds may result from mixed phenotypes as already detected for ciclopirox. Indeed, the profiles of ML228 (1 µM) and SC144 (2 µM) display high similarity to the DNA synthesis cluster that is attributed to their iron chelating properties (Figure 2B) (Schneidewind et al., 2020).

**Figure 2:**
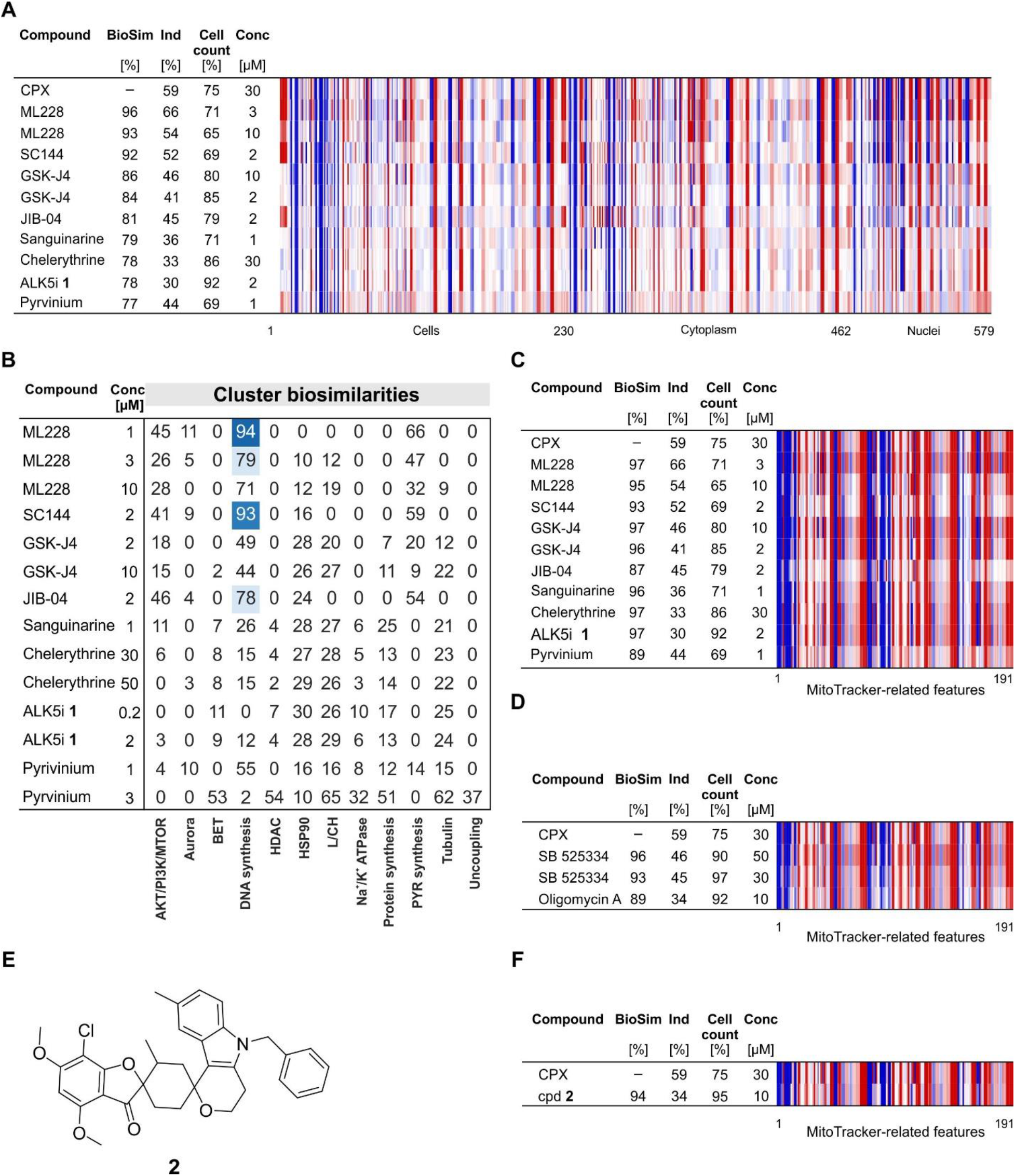
Selected profiles of reference compounds with high biosimilarity to the ciclopirox profile ciclopirox at 30 µM. (A) Morphological profiles of selected reference compounds with biosimilarity ≥ 75 % to the profile of ciclopirox at 30 µM. (B) Cluster biosimilarity heatmap for the profiles of compounds that are biosimilar to ciclopirox at 30 µM. Percent values are given. (C and D) Biosimilarity of the profiles of reference compounds to the profile of ciclopirox at 30 µM when only MitoTracker-related features are considered. (E) Structure of compound **2**. (F) Biosimilarity of the profile of compound **2** (cpd **2**) to the profile of 30 µM ciclopirox when only MitoTracker-related features are considered. (A, C, D and F) The top line of the heatmap profile is set as a reference profile (100 % biological similarity, BioSim) to which the following profiles are compared. Blue color, decreased feature; red color, increased feature. Cpd: compound; BioSim: biosimilarity; Ind: induction; Conc: concentration; L/CH: Lysosmotropism/cholesterol homeostasis; PYR: pyrimidine (see also Figure S2).

We then compared all features related to one of the CPA stains. The biosimilarity of the profiles of these compounds to the ciclopirox profile recorded at 30 µM increased when the MitoTracker-related features were considered, pointing towards an influence of these compounds on mitochondria (see Figure 2C and Figure S2). To only focus on the mitochondrial phenotype, we used all MitoTracker-related features to search for small molecules that share similarity to the profile of ciclopirox at 30 µM and are expected to impair mitochondrial morphology in a similar way. We detected biosimilarity higher than 85 % to the MitoTracker-related features for profiles of more than 20 compounds (see Table S3; of note, as only 191 features were compared, we used the more stringent threshold of 85 % to judge biosimilarity). The profiles of the compounds displayed again high biosimilarity to the ciclopirox profile at 30 µM (Figure 2C). Moreover, the profiles of the ALK5 inhibitor SB525334 and the F_0_F_1_ ATPase inhibitor oligomycin A were biosimilar to the profile of ciclopirox only when the MitoTracker-related features were compared (Figure 2D).

The F_0_F_1_ ATP synthase is part of the mitochondrial electron transport chain (ETC) and uses the mitochondrial proton gradient to generate ATP. We analyzed the profiles of reference compounds that impair ETC such as the complex I inhibitors rotenone, aumitin (Robke et al., 2018), BAY-179 (Mowat et al., 2022), authipyrin (Kaiser et al., 2019) and IACS-010759, the complex II inhibitor lonidamine, complex III inhibitors myxothiazol and antimycin A and the uncoupling agent FCCP. No profile cross-similarity was observed for complex I inhibitors (Figure S3A). Besides targeting complex I, rotenone impairs microtubules (Marshall and Himes, 1978; Srivastava and Panda, 2007) and in CPA, the profile of rotenone is assigned to the tubulin cluster (Akbarzadeh et al., 2022). The profile of aumitin shows similarity to the L/CH cluster at 10 to 50 µM, whereas the profiles of BAY-179 and authipyrin have only low induction values of 6 % at 5 and 10 µM, respectively. Therefore, the detection of complex I activity in CPA remains elusive. As recently reported, the profiles of inhibitors of dihydroorotate dehydrogenase (DHODH, and *de novo* pyrimidine biosynthesis in general) form a CPA cluster with the profiles of complex III modulators as the activity of DHODH is tightly coupled to complex III (Scholermann et al., 2022). No similarity to the ciclopirox profile was detected for the profiles of inhibitors of ETC and the uncoupling agent FCCP using the full CPA profiles (Figure S3B). Using only MitoTracker-related features, biosimilarity to the ciclopirox profile at 30 µM was detected only for the profile of oligomycin A (Figure 2D and Figure S3C). Hence, the detected phenotype is not related to direct impairment of the ETC.

### Analysis of mitochondrial function

To explore the phenotype induced by 30 µM ciclopirox in more detail, GSK-J4 was selected as a second compound with metal ion-chelating properties as well as SB525334 as the profiles of both compounds do not display biosimilarity to the DNA synthesis cluster (Figure 2B and S4A). In the further analysis, the in-house compound **2** was included as an uncharacterized small molecule since its profile was biosimilar to the ciclopirox profile at 30 µM but did not show any similarity to the twelve clusters (Figure 2E-2F and Figure S4B-S4C) (Burhop et al., 2021).

Mitochondria are dynamic organelles that undergo highly coordinated processes of fusion and fission and can rapidly change their shape and function according to the physiological needs of the cells (Scott and Youle, 2010; Willems et al., 2015). Fusion results in the generation of mitochondria that are interconnected and these are present in metabolically active cells (Westermann, 2010). Fission results in numerous mitochondrial fragments and mediates removal of damaged mitochondria (Westermann, 2010). Close inspection of the MitoTracker images revealed altered mitochondrial morphology upon treatment with ciclopirox at 30 and 50 µM with a puncta-like staining that may resemble mitochondrial fragmentation (Figure 3A). A similar pattern was observed for GSK-J4, SB525334 and compound **2** (Figure 3B). To get insight into the phenotype, the mitochondrial network was monitored for 24 h using CellLight™ Mitochondria-GFP BacMam 2.0. In DMSO samples, mitochondria formed elongated structures (see Figure 3C and Movie S1). After treatment with ciclopirox for 10 h, dose-dependent fragmentation of the mitochondrial network became visible, whereas this phenotype evolved faster upon addition of GSK-J4 (Figure 3A, 3C and 3D, and Movie S2-S3). Mitochondrial fragmentation was detected also after treatment with SB525334 and compound **2** (Figure 3A and 3C).

**Figure 3.**
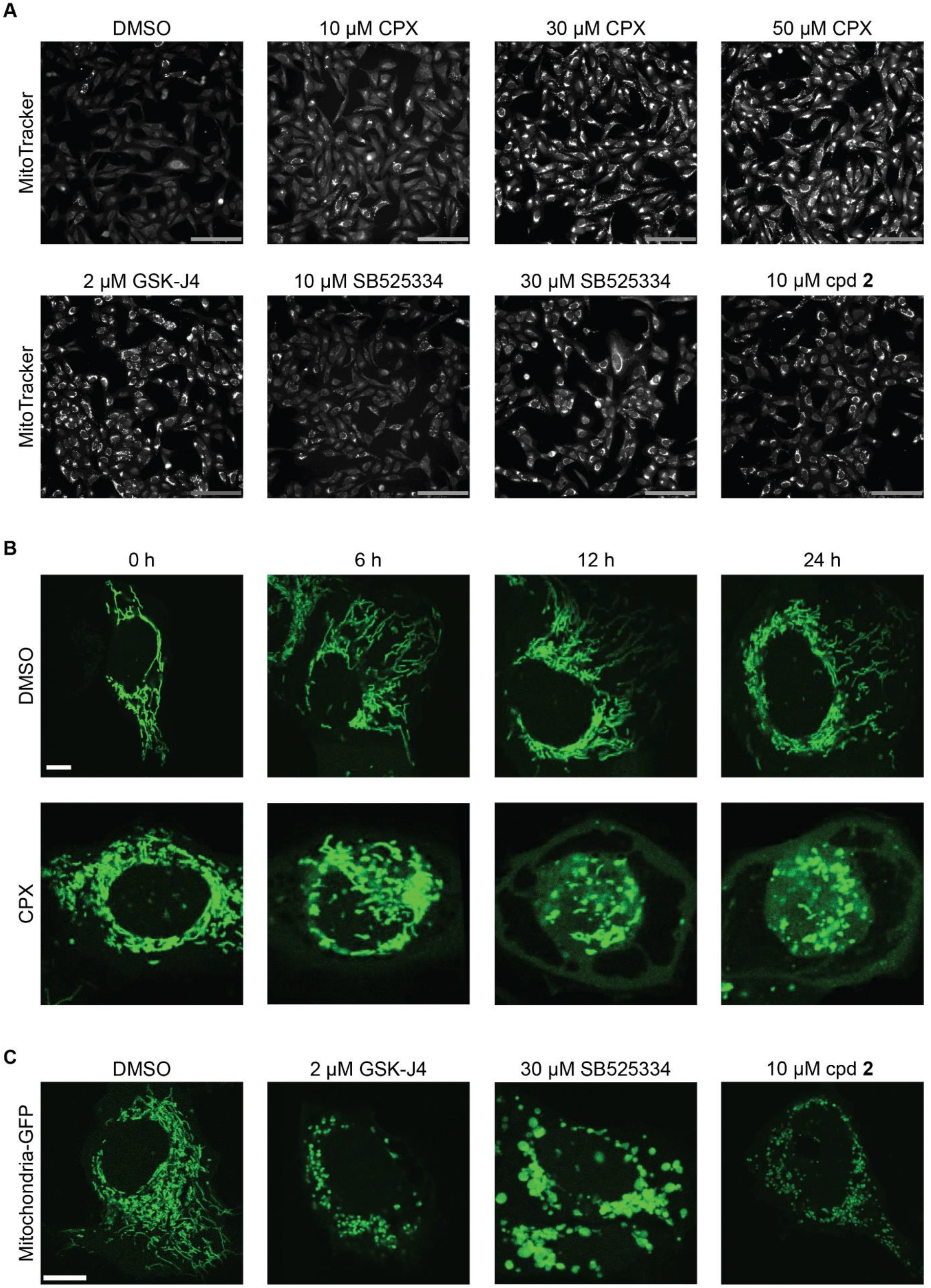
Influence of the compounds on mitochondrial network. (A) MitoTracker Deep Red staining for ciclopirox (CPX), GSK-J4, SB525334 and compound **2** (cpd **2**). Images from CPA are shown. Scale bar: 176 µm. (B and C) U-2OS cells were transduced with CellLight^TM^ Mitochondria-GFP BacMam 2.0 and treated with the compounds. (B) Mitochondrial morphology was observed in real-time over 24 h after treatment with 30 µM ciclopirox. Images at selected time points are displayed. Scale bar: 10 µm. (C) Mitochondrial morphology after treatment with GSK-J4, SB525334 and compound **2** for 24 h compound. Scale bar: 10 µm. See also Movies S1-S3.

Mitochondria are involved in the induction of apoptosis and mitochondrial fragmentation occurs early during apoptotic process (Arnoult, 2007) and therefore the detected phenotype may be related to cell death. The influence on cell growth and cell death was analyzed by real-time live-cell imaging in U-2OS cells over 48 h. Propidium iodide (PI) and caspase 3/7 activity were used as markers of cell toxicity and apoptosis, respectively (Figure S5). Cell growth was hardly impaired by the compounds even after 48 h and only a slight decrease in cell confluence was detected for 10 µM GSK-J4 (Figure S5). Therefore, the observed mitochondrial phenotype is not related to cell death.

The influence on mitochondrial respiration by means of metabolic flux analyses was assessed using the Seahorse technology and determined the oxygen consumption rate (OCR) and extracellular acidification rate (ECAR) as a measure of mitochondrial respiration and glycolysis, respectively (Figure 4A-4C). Hardly any influence on OCR and ECAR was observed after acute injection of ciclopirox, GSK-J4, SB525334 and compound **2** to U-2OS cells (Figure 4A and 4C). However, dose-dependent suppression of oxygen production was detected after 24 h treatment with ciclopirox and GSK-J4, which was almost completely suppressed at 30 µM ciclopirox and 10 µM GSK-J4 (Figure 4B). Similar results were obtained for SB525334 and compound **2** (Figure 4B and 4D). The drop in OCR may result in an increase of ECAR, which occurs as a compensatory flux (Mookerjee et al., 2016). Indeed, increased extracellular acidification was detected for all compounds at the highest tested concentrations (Figure 4B-4D). These findings point towards suppression of mitochondrial respiration by the compounds that evolves over time.

**Figure 4.**
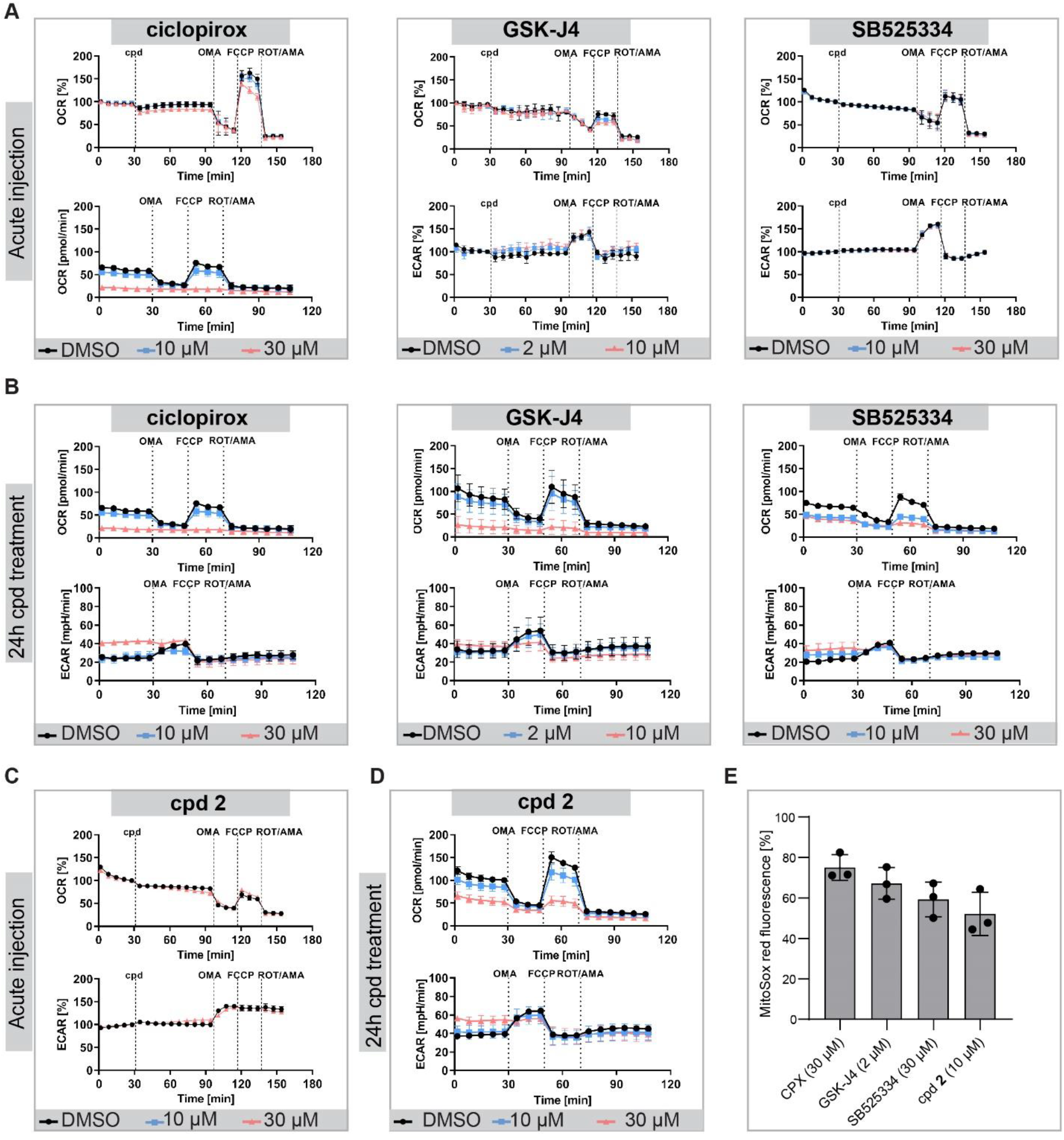
Influence of the compounds on mitochondrial function. (A-C) Influence on mitochondrial respiration as determined using the Seahorse technology and a MitoStressTest. The oxygen consumption rate (OCR) and extracellular acidification rate (ECAR) were measured using the Seahorse XF_p_ analyzer. (A and C) Acute injection of ciclopirox, GSK-J4 and SB525334, and compound **2** (cpd **2**) to U-2OS cells. (B and D) U-2OS cells were treated with the compounds for 24 h prior to measuring OCR and ECAR. Mean values ± SD (n = 3). OMA: oligomycin A; ROT: rotenone; AMA: antimycin A. (E) Detection of mitochondrial superoxide using MitoSOX Red after 1 h treatment with the compounds. Data are normalized to CDNB (positive control, set to 100 %) and DMSO (negative control, set to 0 %). Mean values ± SD (n = 3).

Loss of mitochondrial membrane potential or increased production of reactive oxygen species (ROS) can lead to mitochondrial fragmentation (van der Bliek et al., 2013; Willems et al., 2015). The influence of the compounds on the mitochondrial membrane potential was assessed using TMRE (tetramethylrhodamine, ethyl ester) as a marker (Zorova et al., 2018). TMRE freely crosses membranes regardless of the potential and localizes to active mitochondria due to its negative charge, while it cannot accumulate in depolarized, i.e., dysfunctional mitochondria (Scaduto and Grotyohann, 1999). After 24 h incubation of cells with the compounds, no notable loss of membrane potential compared to the controls could be discerned (Figure S6), while FCCP showed a pronounced reduction of TMRE intensity that is in line with its uncoupling activity (Park et al., 2002). Furthermore, all four compounds increased the level of mitochondrial superoxide as detected with the fluorogenic indicator MitoSox Red (Robinson et al., 2006) which indicates the induction of oxidative stress (Figure 4E).

#### Proteome-wide analysis upon compound perturbation

To gain insight into the modulated pathways, we explored the difference in the proteomes upon treatment of U-2OS cells for 24 h with ciclopirox, GSK-J4, SB-525334 and compound **2** (see Figure 5 and 6 and Figure S7). In the presence of ciclopirox, more than 400 proteins were differentially regulated at 30 µM as compared to ca. 200 regulated proteins at 10 µM ciclopirox (see Figure 5A-5C and Table S4). 114 proteins were upregulated and 62 proteins were downregulated at both concentrations (Figure 5C and Table S4 and S5). Pathway overrepresentation analysis of the proteome at 30 µM ciclopirox linked the altered levels of these proteins to modulation of glucose metabolism, glycolysis, oxidative phosphorylation (OXPHOS), mitochondrial dysfunction and hypoxia-inducible factor 1 (HIF1) signaling (Figure 5D).

**Figure 5.**
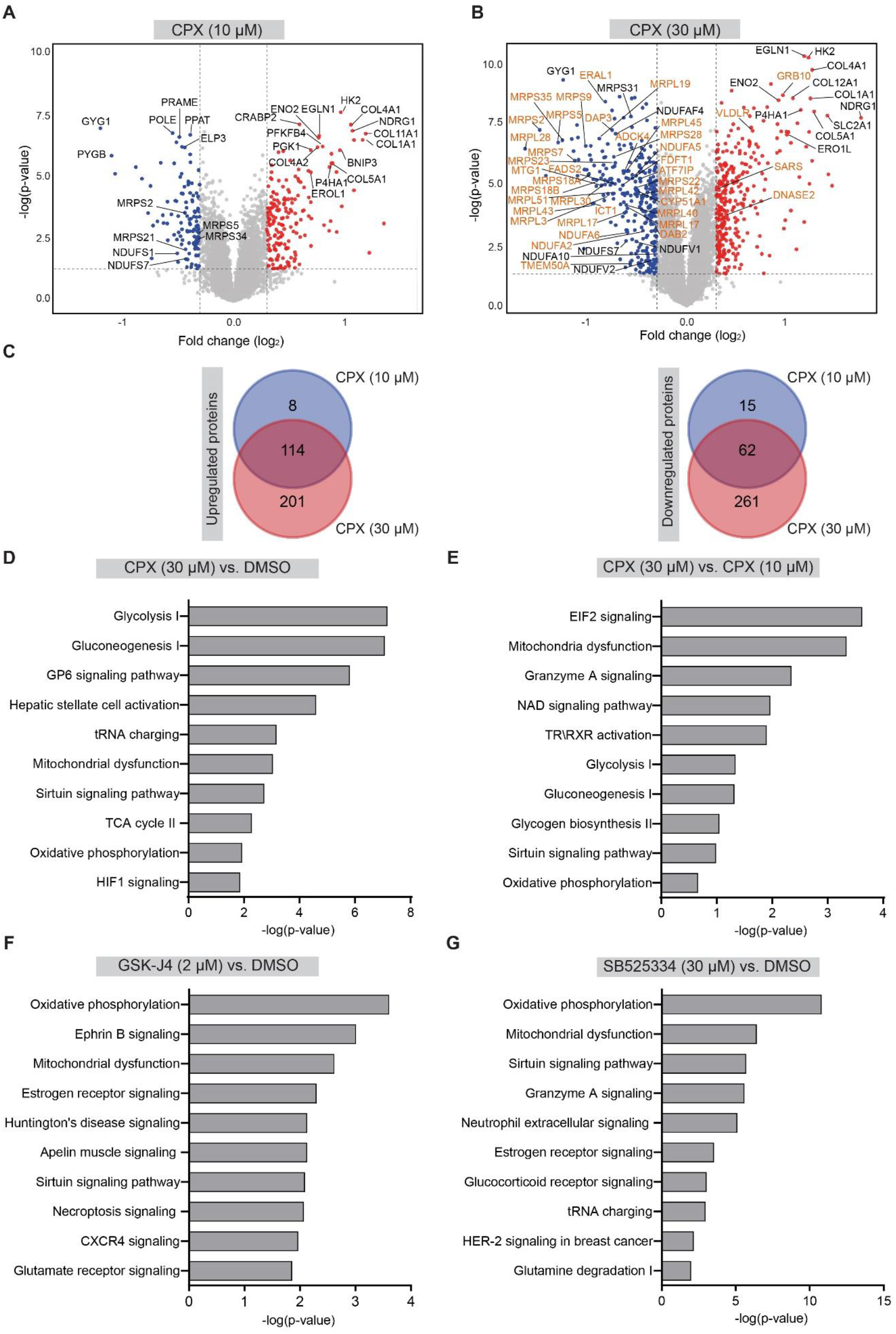
Global proteome profiling and pathway enrichment analysis. (A and B) Volcano plot of log2 fold changes for ciclopirox at 10 and 30 μM upon 24 h compound treatment. Red circles: upregulated proteins; blue circles: downregulated proteins; orange circles: proteins found regulated in Quiros et al. Volcano plots were visualized using VolcaNoseR (Goedhart and Luijsterburg, 2020) (C) Comparison of the number of upregulated and downregulated proteins by 30 µM vs. 10 µM ciclopirox. See also Table S4. (D-E) Pathway overrepresentation analysis for 30 µM ciclopirox in comparison to DMSO (D) or to 10 µM ciclopirox (E). (F and G) Pathway overrepresentation analysis for or 2 µM GSK-J4 (F) or 30 µM SB525334 (G) compared to the DMSO control. TCA: tricarboxylic acid cycle (see also Figure S7 and Tables S5 and S6).

**Figure 6.**
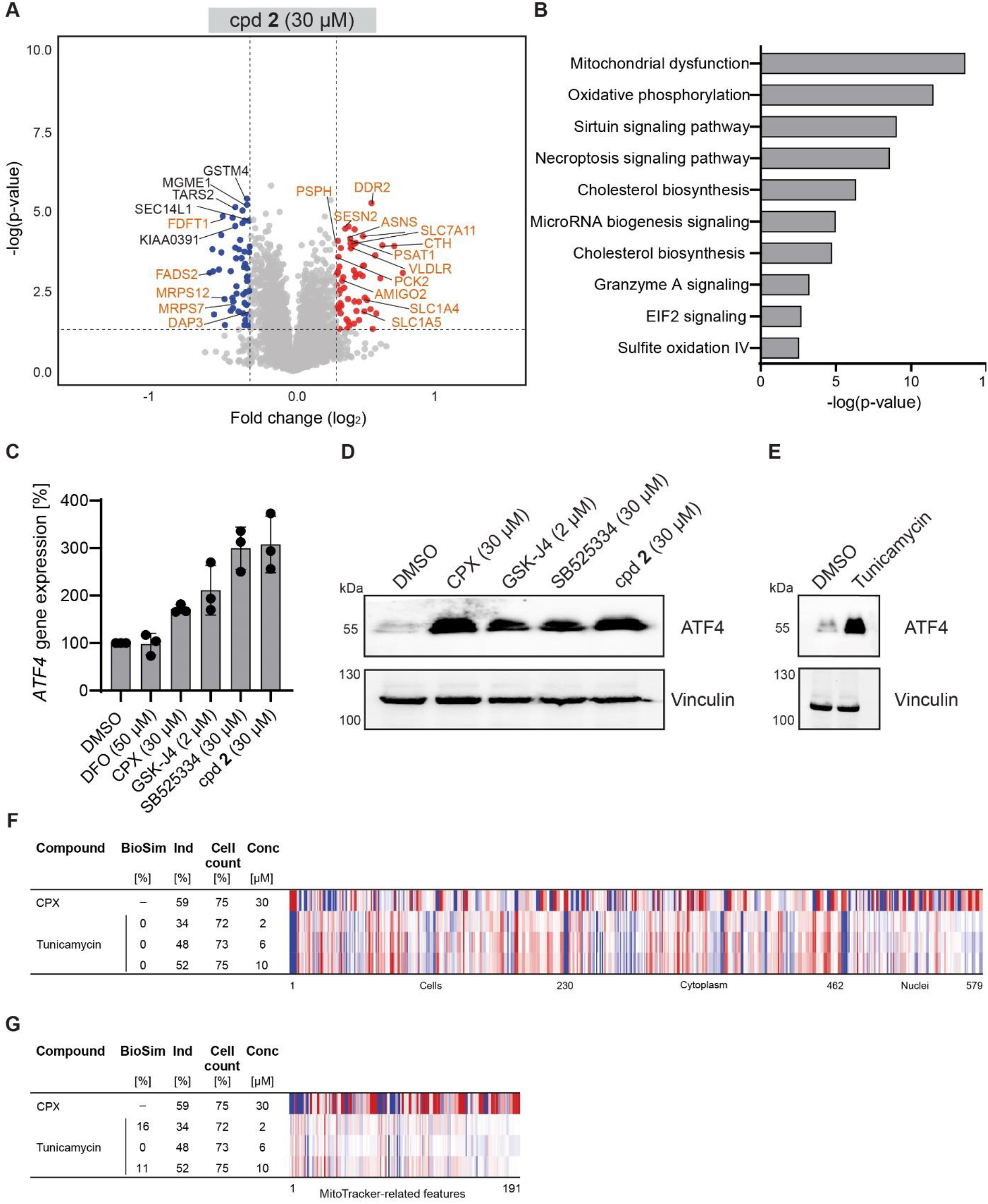
Global proteome profiling for compound 2 and ATF4 regulation. (A) Volcano plot of log2 fold changes for 30 µM compound **2** upon 24 h compound treatment. Red circles: upregulated proteins; blue circles: downregulated proteins. orange circles: proteins found regulated in Quiros et al. Volcano plot was visualized using VolcaNoseR (Goedhart and Luijsterburg, 2020) (B) Pathway overrepresentation analysis for compound **2**. See also Table S6. (C) Upregulation of *ATF4* gene expression upon compound treatment. U-2OS cells were treated with the compounds for 24 h prior to detection of ATF4 mRNA levels using RT-qPCR. Mean values ± SD (n =3). (D and E) Detection of ATF4 protein levels. U-2OS cells were treated with the compounds for 24 h prior to detection of ATF4 and vinculin as a reference protein using immunoblotting. Tunicamycin: 1 µg/ml. Representative blot is shown (n =3). See Figure S9 for the full blots. (F and G) Comparison of the profiles of tunicamycin to the ciclopirox profile at 30 µM. (F) Full profiles. (G) Comparison of MitoTracker-related features. The top line profile is set as a reference profile (100 % biological similarity, BioSim) to which the following profiles are compared. Blue color: decreased feature; red color: increased feature. Cpd: compound; BioSim: biosimilarity; Ind: induction; Conc: concentration.

Glucose metabolism and HIF1 signaling are linked since during hypoxia, cells switch their metabolism from mitochondrial respiration to glycolysis to meet the bioenergetic requirements (Kierans and Taylor, 2021). Ciclopirox upregulates several proteins such as the HIF1 prolyl hydroxylase Egl nine homolog 1 (EGLN1), hexokinase-2 (HK2), heme oxygenase 1 (HMOX1) and the glucose transporter solute carrier family 2 member 1 (SLC2A1, also known as GLUT-1) (Table S4-S6). These proteins are involved in HIF1 signaling or their expression is induced by the HIF1 transcription factor. The HIF1-α protein is regulated on the protein level as prolyl hydroxylation of HIF1-α by HIF prolyl hydrohylase (HIF PHD) leads to its proteasomal degradation via the E3 ligase Von Hippel-Lindau (VHL). As HIF PHD requires oxygen for its enzymatic activity, HIF1-α levels are low during normoxia and increase during hypoxia. Furthermore, HIF PHD require Fe(II) as a cofactor. Therefore, iron chelators induce HIF1 response by interfering with the activity of prolyl hydroxylases, thereby stabilizing HIF1 (Kaelin and Ratcliffe, 2008). Ciclopirox has been reported to stabilize HIF1-α under normoxic conditions (Linden et al., 2003; Milosevic et al., 2009). We therefore explored whether compounds biosimilar to 30 µM ciclopirox also regulate HIF1 levels. No modulation of HIF1 signaling was observed in the proteome profiling for GSK-J4, SB525334 and compound **2** (Figure 5F and 5G, Figure S7 and Figure 6A and B). Moreover, the compounds neither induced HIF1-α -dependent reporter expression nor increased HIF1 protein levels (Figure S8A and S8B). In line with this, only ciclopirox scored as a direct iron chelator using ferrozine-base iron chelation assay (Figure S8C). Hence, modulation of HIF1 signaling is not the common denominator for the detected phenotype in CPA.

Comparison of the proteomes at 30 µM and 10 µM ciclopirox revealed eukaryotic translation initiation factor 2 (EIF2) signaling as the most significantly modulated process at 30 µM ciclopirox (Figure 5E). Among the downregulated proteins upon treatment with 30 µM ciclopirox vs. 10 µM ciclopirox were several mitochondria ribosomal proteins (MRPs) and NADH-ubiquinone oxidoreductase subunits (NDUFs), which are complex I components (Figure 5A and 5B, Table S4). The most significantly regulated pathways for all tested compounds were oxidative phosphorylation and mitochondrial dysfunction (Figure 5F and 5G and Figure 6A-6B). Modulation of MRPs and/or NDUFS by small molecules has been observed for small molecules that induce mitochondrial stress response such as FCCP, doxycycline, actinonin and MitoBLoCK-6 and these changes were only mapped on proteome but not transcriptome level (Quirós et al., 2017). In parallel, these compounds induced the expression of genes that are regulated by cAMP dependent transcription factor 4 (also known as activating transcription factor, ATF4) (Quirós et al., 2017). ATF4 expression is repressed under normal conditions and induction of ATF4 is a hallmark of the integrated stress response (ISR) (Neill and Masson, 2023). ISR is a signaling pathway that is activated as a response of altered physiological conditions such as endoplasmic stress, hypoxia, glucose or amino acid deprivation or viral infections, which all lead to phosphorylation of EIF2α (Pakos-Zebrucka et al., 2016). As a consequence, global protein translation is reduced, while ATF4 expression helps the cell to recover and survive (Pakos-Zebrucka et al., 2016). The extent and duration of ISR determines cell fate and may also lead to cell death.

Several proteins that were found downregulated by Quiros et al. were also downregulated by ciclopirox, SB525334 and compound **2** but not by GSK-J4. Proteins regulated by 30 µM ciclopirox displayed the highest overlap (see Table S7). Moreover, some of the genes or proteins that were reported as upregulated by Quiros et al. were present also at higher levels upon treatment with ciclopirox, GSK-J4 and compound **2** and compound **2** showed the highest overlap (Table S8). These findings suggested that the compounds may induce ATF4 transcriptional response and ISR. Therefore, we explored whether the compounds affect ATF4 expression. Whereas no ATF4 mRNA and protein was detected in the control condition, ciclopirox, GSK-J4, SB525334 and compound **2** increased the *ATF4* gene expression (Figure 6C). In line with this result, ATF4 protein was detected upon treatment with all four compounds (Figure 6D), indicating activation of the integrated stress response. In contrast, the iron chelator DFO, whose CPA profiles are not biosimilar to 30 µM ciclopirox, does not stimulate *ATF4* expression (Figure 6C). Tunicamycin is a natural product that decreases N-glycosylation in cells and induces endoplasmic stress and ATF4 expression (Luhr et al., 2019). We detected increased levels of ATF4 protein upon treatment of U-2OS cells with tunicamycin (Figure 6E). However, the CPA profiles of tunicamycin were not biosimilar to the profile of 30 µM ciclopirox, both, using the full profiles and only MitoTracker-related features (Figure 6F and 6G). Hence, CPA can distinguish between endoplasmic and mitochondrial stress/ISR, even though both operate via induction of ATF4. These findings reveal the induction of mitochondrial stress and ISR as the common MoA for compounds sharing the same morphological changes as ciclopirox at 30 µM.

### Definition of a cluster related to mitochondrial stress

To enable fast prediction of the observed biological activity, we used the CPA profiles of selected compounds that are biosimilar to the profile recorded in the presence of 30 µM ciclopirox (see Table S9) to extract the characteristic CPA features for this cluster (termed MitoStress cluster) and to obtain a cluster subprofile according to the recently described procedure (Pahl et al., 2023). The MitoStress cluster subprofile consists of 294 features and is very different from the twelve defined clusters thus far as demonstrated by the cluster profile cross-correlation and the lower dimension UMAP plot (Figure 7A-7B and Figure S10). Thus, the MitoStress cluster is a very valuable new cluster for analysis of morphological profiling by means of CPA.

**Figure 7.**
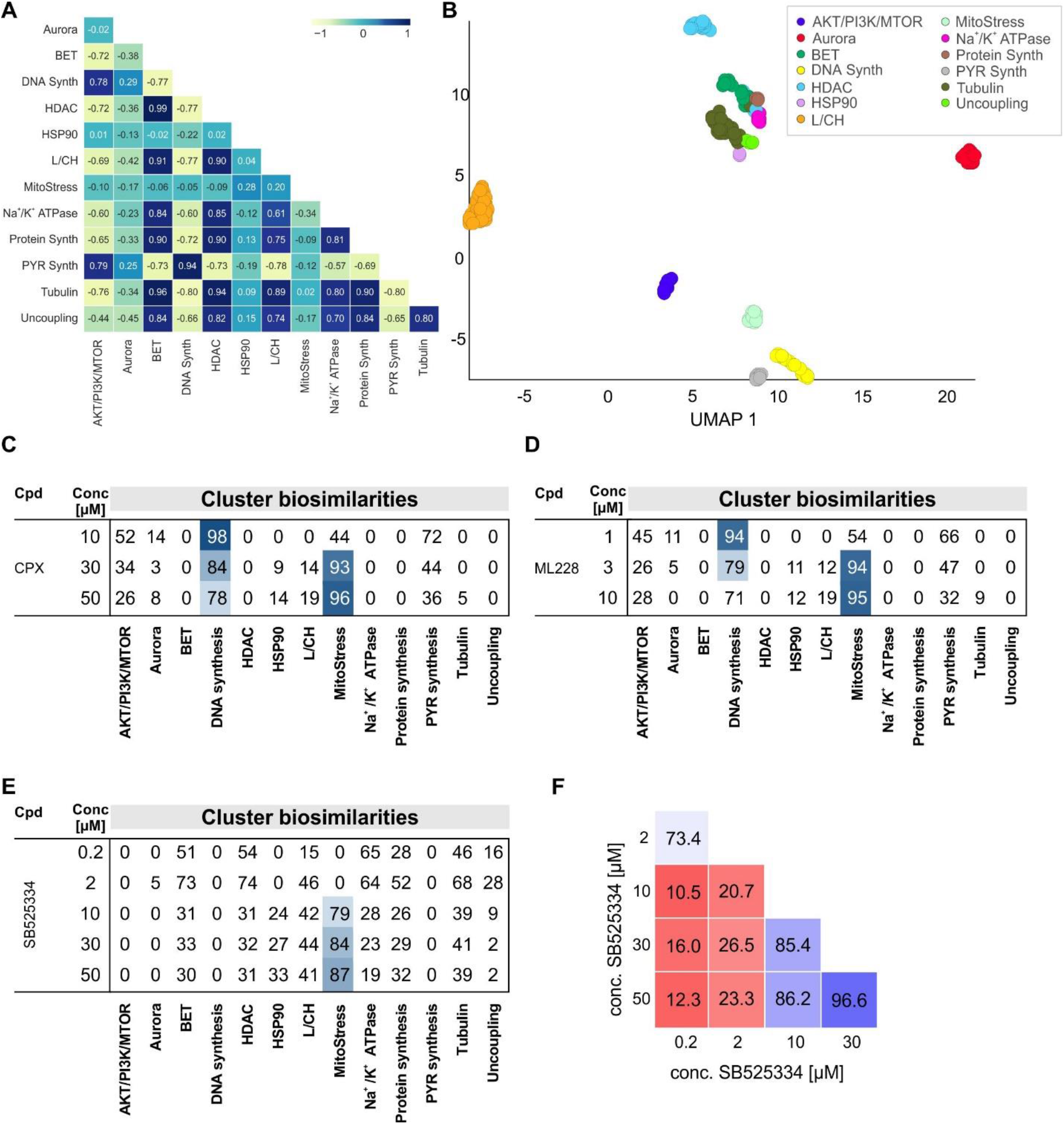
Definition of the MitoStress cluster. (A) Cluster subprofile cross-correlation using Pearson correlation. (B) UMAP plot using the full profiles of the reference compounds that were used to define the bioactivity clusters. Not normalized data, 10 neighbors. (C-E) Cluster biosimilarity heatmap for the profiles of ciclopirox (CPX) (C), ML228 (D) and SB525334 (E). Profile cross-correlation for SB525334. Values are biosimilarity in %. L/CH: Lysosomotropism/cholesterol homeostasis; PYR: pyrimidine. Synth: synthesis.

A cluster biosimilarity map clearly depicts the morphological phenotypes caused by ciclopirox: whereas at 10 µM profile similarity is observed only to the DNA synthesis cluster, additional similarity to the MitoStress cluster is detected for the ciclopirox profile at 30 µM, while the similarity to the DNA synthesis cluster decreases (Figure 7C). These morphological changes are dose dependent and more pronounced at 50 µM (Figure 7C). A similar ‘cluster shift’ was observed for the profile of the iron chelator ML228 (Figure 7D) but not for the profiles of DFO, phenanthroline, deferasirox and PAC-1 (Figure S11A-S11D). Hence, all tested iron chelating compounds influence DNA synthesis but not all of them impair mitochondrial function in the tested concentration range. High similarity to the MitoStress cluster is detected for the profiles of further cluster members (see Figure S11E). For the ALK5 inhibitor SB525334, the MitoStress phenotype starts evolving already at 10 µM and the MitoStress cluster similarity increases in a concentration-dependent manner (Figure 7E). Importantly, for several compounds the MitoStress phenotype is detected at higher concentrations as exemplified by ciclopirox, ML228, SB525334, Sal003 and the kinase inhibitor kenpaullone (Figure 7C-7E and Figure S11F and S11G). A cluster shift may result in less biosimilar or even dissimilar profiles at different concentrations of a given compound as detected for SB525334, Sal003 and kenpaullone (see Figure 7F and Figure S11H and S11I). The biosimilarity to the MitoStress cluster for all CPA-active reference compounds, which we have profiled thus far, can be assessed using the web app tool https://cpcse.pythonanywhere.com/.

## Discussion and Conclusions

Detailed knowledge about the targets and processes impaired by small molecules is essential for their proper use as research tools and, more importantly, for drug candidates. Small molecules are usually identified in assays monitoring modulation of a target or a process, either *in vitro*, in cells or *in vivo*. Thorough characterization of bioactive compounds informs about efficacy and selectivity and, in addition, safety-panel profiling can uncover off-target liabilities. Off-and on-target adverse effects are best identified early in the compound development workflow and profiling approaches in cells such as transcriptomics, proteomics and morphological profiling provide an unbiased view on various bioactivities of small molecules. CPA determines morphological profiles upon perturbation (Ziegler et al., 2021) and, in principle, comparison to the profiles of annotated compounds should lead to target hypotheses. In fact, often similarity in CPA profiles is not linked to the same target annotation, which may result from off-target activity or impairment of the same pathway but at a different level (Akbarzadeh et al., 2022; Schneidewind et al., 2020; Schneidewind et al., 2021; Scholermann et al., 2022).

Analyzing the CPA profiles of reference compounds revealed a new CPA bioactivity cluster that impairs mitochondrial morphology by inducing a fragmented phenotype and mitochondrial stress. These morphological changes were detected for some but not all iron-chelating compounds. Whereas ciclopirox, ML228, GSK-J4 and SC144 induced mitochondrial stress at the tested concentrations, DFO, phenanthroline, deferasirox and PAC-1 did not. Induction of HIF1 by the chelator desferrioxamine causes mitochondrial fission (Marsboom et al., 2012). However, while all tested iron chelators inhibit DNA synthesis, they do not share mitochondrial fission and stress as a common MoA. The detected differences for iron chelators underscore the power of morphological profiling in mapping various bioactivities for given compounds, which assists selection and proper use of these small molecules in cellular studies. Of note, Wheeler et al. identified ciclopirox, ML228, GSK-J4, JIB-04, SC144 and NSC319726 as hits in a screen for macrofilaricidal activity in *C. elegans* (Wheeler et al., 2022) and the activity could not be linked to the annotated targets (e.g., histone demethylases, HIF-1α). The macrofilaricidal activity may be due to iron chelation. Alterative MoA is suggested by our CPA study and may link mitochondrial stress to macrofilaricidal activity.

CPA has been employed to study cell and mitochondrial toxicity (Dahlin et al., 2023; Garcia de Lomana et al., 2023; Herman et al., 2023; Seal et al., 2022; Trapotsi et al., 2022). By using CPA profiles, gene expression signatures, chemical structural information and mitochondrial toxicity data, Seal *et al*. could distinguish between mitochondrial toxicants and non-toxicants based on the morphological profiles (Seal et al., 2022). None of the CPA profiles of the mitotoxic compounds investigated by Seal *et al*. display similarity to the MitoStress cluster (Figure S12A) and, therefore, impair mitochondrial homeostasis by a different MoA. Trapotsi *et al*. explored protein-targeting chimeras in CPA along with mitotoxic compounds to train a model for mitotoxicity (Trapotsi et al., 2022). The CPA profiles for the disclosed compounds are not biosimilar to the MitoStress cluster (Figure S12B). In line with this, we also did not detect similarity to the MitoStress cluster for compounds directly impairing the mitochondrial ETC. Moreover, the MitoStress phenotype, which we detect after treatment for 20 h, is not linked to toxicity as we failed detecting cell death even 48 h after compound addition. Hence, we identified a bioactivity cluster linked to mitochondrial stress that can be mapped using morphological profiling.

The compounds in this cluster cause profound changes in the mitochondrial network architecture that is reminiscent of mitochondrial fragmentation. Several compounds have been reported as fission inducers (Zorov et al., 2019). The iron chelator phenanthroline induces mitochondrial fragmentation at 50 µM (Park et al., 2012) and also we detect biosimilarity between the profiles of 30 µM ciclopirox and 50 µM phenanthroline. However, the similarity of the profile of phenanthroline at 50 µM to the MitoStress cluster (62 %) is lower than the suggested biosimilarity threshold of 80 % for subprofile comparison. In general, the similarity to the MitoStress cluster is detected at higher concentrations, whereas the phenotype differs at lower concentrations as observed for the iron chelators ciclopirox and ML228 but also for SB525334, Sal003 and kenpaullone. Therefore, the morphological profiles for this type of compounds can guide the selection of an appropriate concentration for cellular experiments in order to study the desired mechanism of action without causing mitochondrial stress. This is particularly important considering that the MitoStress phenotype occurs at non-toxic concentrations and may be therefore easily overlooked.

Mitochondrial fragmentation is linked to oxidative stress (Hung et al., 2018; Willems et al., 2015) and we detected increased levels of mitochondrial superoxide by ciclopirox, GSK-J4, SB525334 and compound **2**. The compounds did not attenuate the mitochondrial membrane potential and suppressed mitochondrial respiration only after long-term treatment. ETC dysfunction is linked to mitochondrial fission (Liu et al., 2022). Mitochondrial fragmentation has been reported for ETC inhibitors and uncoupling agents but for them, we did not detect the fission phenotype in U-2OS cells and biosimilarity to the MitoStress cluster at the tested concentrations after treatment for 20 h. Hence, CPA can differentiate between different types of mitochondrial modulators such as complex III inhibitors (which are assigned to the pyrimidine synthesis cluster), uncoupling agents and compounds inducing mitochondrial stress.

Besides the four small molecules explored here, the profiles of further reference compounds displayed biosimilarity to the profile recorded for ciclopirox at 30 µM and the MitoStress cluster. Some of them are lipophilic cationic molecules that have been reported to influence mitochondria and may accumulate in these organelles (Lei et al., 2018). For example, the alkaloids sanguinarine and chelerythrine increased ROS levels and attenuate mitochondrial membrane potential (Mick et al., 2020). Pyrvinium pamoate inhibited mitochondrial respiration and induced ISR in MOLM13 cells (Fu et al., 2021). The phosphatase inhibitor Sal003 is an inhibitor of the EIF2α phosphatase and thereby promotes EIF2 phosphorylation that ultimately increases ATF4 levels, thus protecting cells from endoplasmic reticulum stress (Boyce et al., 2005; Costa-Mattioli et al., 2007). We detected similarity of Sal003 to the MitoStress cluster at 10 µM but not at 3 or 6 µM. For its less potent derivative salubrinal, neither similarity to the MitoStress cluster is detected nor profile similarity to Sal003 up to concentrations of 50 µM (Figure S13). The low induction values for salubrinal in comparison to Sal003 (see Figure S13B and S13C) are in line with its lower potency and explain the observed differences in CPA.

Treatment with 30 µM ciclopirox reduced the levels of numerous MRP and NDUF proteins that indicates downregulation of mitochondrial translation and complex I activity. Quiros et al. observed lower levels for several MRPs and NDUFs in a study using mitochondrial stressors such as doxycycline, FCCP, actinonin and MitoBlock 6 (Quirós et al., 2017). This regulation occurs on translation level as no changes in the expression of the corresponding coding genes were detected. Increase in ROS production can cause global inhibition of protein synthesis by different mechanisms and leads to phosphorylation and inactivation of EIF2 (Grant, 2011) (Samluk et al., 2019). At the same time, mRNAs with upstream open reading frame (uORF) are selectively translated such as for the transcription factor ATF4 (Jenkins et al., 2021). ATF4 activates the transcription of genes involved in amino acid transport, serine biosynthesis, one carbon metabolism, antioxidant defense and proteostasis (Liu et al., 2022; Mick et al., 2020), a stress pathway known as integrated stress response (ISR). In line with this, Quiros et al. observed upregulation of genes involved in serine biosynthesis and one carbon metabolism, and demonstrated the activation of ATF4 and ISR (Quirós et al., 2017). Similarly, we detected elevated levels of *ATF4* mRNA and ATF4 protein by the four studied compounds, indicating that the observed CPA phenotype is due to oxidative stress, mitochondrial fragmentation and ISR. This is supported by the fact that the iron chelator DFO fails inducing mitochondrial fragmentation and ATF4 expression and, therefore, this phenotype is not caused by general iron chelation.

Mitochondrial stress duration has a different impact on ATF4 levels: whereas ATF4 is induced after short-term stress, ATF4 signaling is attenuated after long-term stress to allow for protection against long-lasting inhibition of protein synthesis (Samluk et al., 2019). This finding together with the type and potency of the mitochondrial stressor may explain why some known mitochondrial stressor such as FCCP or antimycin A lead to a different CPA phenotype: FCCP shares a distinct phenotype with other, structurally dissimilar uncouplers, whereas the profile of antimycin A is assigned to the pyrimidine synthesis cluster (Pahl et al., 2023; Scholermann et al., 2022).

The extent and duration of ISR determines the cell fate and may lead to cell death. Several compounds share in CPA the mitochondrial fragmentation/ISR phenotype. Considering that ISR is implicated in different diseases (Costa-Mattioli and Walter, 2020), strategies for pharmacological modulation of ISR have been explored (Costa-Mattioli and Walter, 2020; Pakos-Zebrucka et al., 2016). To allow for easy detection of compounds inducing mitochondrial stress and ISR, we defined the MitoStress cluster by extracting a median profile of the observed morphological changes. Biosimilarity to this cluster subprofile would suggest the induction of mitochondrial stress and ISR. For all CPA-active reference compounds that we have screened thus far, biosimilarity to the MitoStress cluster subprofile can be queried using the web app tool https://cpcse.pythonanywhere.com/.

In summary, using CPA we identified a morphological phenotype that is related to mitochondrial stress and is linked to the activation of ATF4 and the integrated stress response. This MoA is shared by compounds with different targets that indirectly suppress mitochondrial respiration and increase ROS levels in mitochondria and differs from the CPA profiles of direct inhibitors of ETC or uncouplers. The newly defined MitoStress cluster and cluster subprofile will enable rapid prediction of this MoA for compounds profiled using CPA.

## Acknowledgments

Research at the Max Planck Institute of Molecular Physiology was supported by the Max Planck Society. This work was co-funded by the European Union (Drug Discovery Hub Dortmund (DDHD), EFRE-0200481) and Innovative Medicines Initiative (grant agreement number 115489) resources of which are composed of financial contribution from the European Union’s Seventh Framework Programme (FP7/2007-2013) and EFPIA companies’ in-kind contribution, and by EPSRC (grants EP/N025652/1 and EP/F043503/1). This work was funded from the programme “Netzwerke 2021”, an initiative of the Ministry of Culture and Science of the State of Northrhine Westphalia. We thank Prof. Dr. Konstanze Winklhofer and Dr. Verian Bader for helpful discussions. The compound management and screening center (COMAS) in Dortmund is acknowledged for performing the high-throughput screening. The HRMS team at the Max Planck Institute is acknowledged for performing the mass spectrometry.

## Author Contributions

S.Z. and H.W. designed the research. S.R.A., D.A., S.K. and J.W. performed biological experiments. A.B. synthesized compounds. S.S. and A.P. performed CPA and processed the data. P.J. and M.M. analysed the proteomics data. S.Z. and S.R.A. analyzed the CPA data. S.R.A., S.Z. and H.W. wrote the manuscript. All authors discussed the results and commented on the manuscript.

