## Supporting Information for "Detection of a Mitochondrial Stress Phenotype using the Cell Painting Assay"

**Supplementary Figure S1-13**

**Tables S1-S9**

**Movies S1-S3**

**Material and Experimental Section**

### Supporting Figures

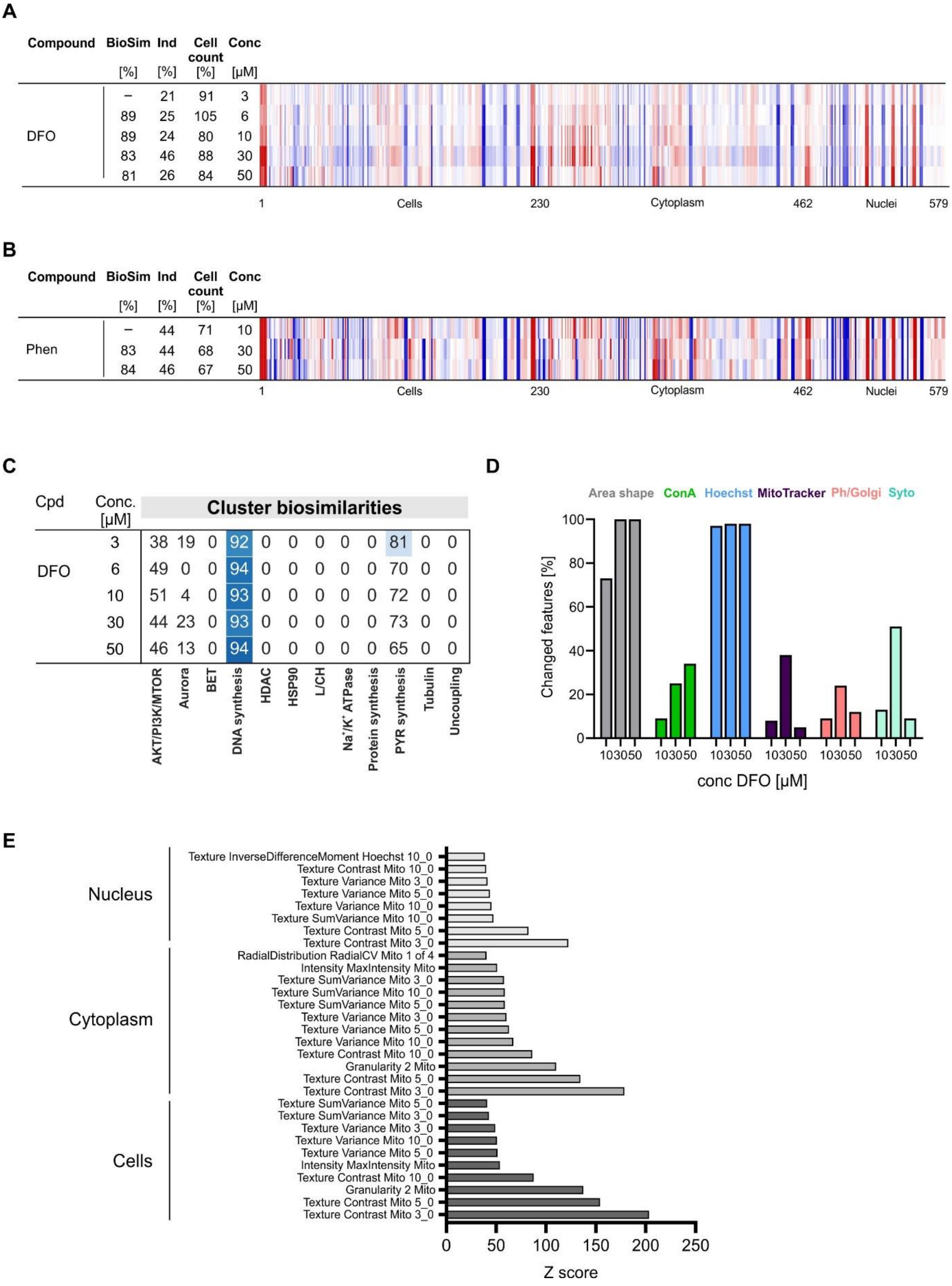

**Figure S1 (related to Figure 1): CPA profiles for deferoxamine (DFO) and phenanthroline.** (A and B) Comparison of the profiles for DFO (A) and phenanthroline (B) at different concentrations. The top line profile is set as a reference profile (100 % biological similarity, BioSim) to which the following profiles are compared. Blue color: decreased feature; red color: increased feature. (C) Cluster biosimilarity heatmap for DFO. Percent values are displayed. (D) Dose-dependent change in dye-related CPA features at different concentrations DFO. (E) Z scores for top 30 altered features with high Z scores for ciclopirox at 30 µM as determined in CPA. Cpd: compound; BioSim: biosimilarity; Ind: induction; Conc: concentration.

| Compound | BioSim [%] | Ind [%] | Cell count [%] | Conc [µM] |
|---|---|---|---|---|
| CPX | – | 59 | 75 | 30 |
| ML288 | 94 | 66 | 71 | 3 |
| ML288 | 91 | 54 | 65 | 10 |
| SC144 | 92 | 52 | 69 | 2 |
| GSK-J4 | 81 | 46 | 80 | 10 |
| GSK-J4 | 79 | 41 | 85 | 2 |
| JIB-04 | 78 | 45 | 79 | 2 |
| Sanguinarine | 70 | 36 | 71 | 1 |
| Chelerythrine | 71 | 33 | 86 | 30 |
| cpd **1** | 68 | 30 | 92 | 2 |
| Pyrvinium | 74 | 44 | 69 | 1 |

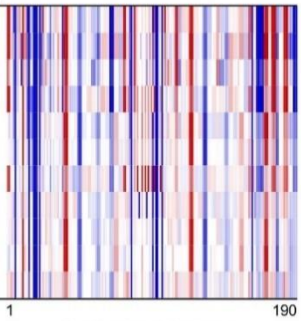


| Compound | BioSim [%] | Ind [%] | Cell count [%] | Conc [µM] |
|---|---|---|---|---|
| CPX | – | 59 | 75 | 30 |
| ML288 | 97 | 66 | 71 | 3 |
| ML288 | 95 | 54 | 65 | 10 |
| SC144 | 94 | 52 | 69 | 2 |
| GSK-J4 | 78 | 46 | 80 | 10 |
| GSK-J4 | 78 | 41 | 85 | 2 |
| JIB-04 | 76 | 45 | 79 | 2 |
| Sanguinarine | 59 | 36 | 71 | 1 |
| Chelerythrine | 54 | 33 | 86 | 30 |
| cpd **1** | 49 | 30 | 92 | 2 |
| Pyrvinium | 68 | 44 | 69 | 1 |

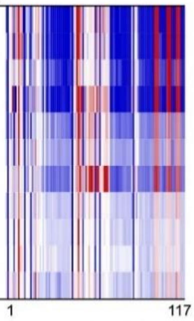


| Compound | BioSim [%] | Ind [%] | Cell count [%] | Conc [µM] |
|---|---|---|---|---|
| CPX | – | 59 | 75 | 30 |
| ML288 | 90 | 66 | 71 | 3 |
| ML288 | 91 | 54 | 65 | 10 |
| SC144 | 85 | 52 | 69 | 2 |
| GSK-J4 | 93 | 46 | 80 | 10 |
| GSK-J4 | 87 | 41 | 85 | 2 |
| JIB-04 | 63 | 45 | 79 | 2 |
| Sanguinarine | 83 | 36 | 71 | 1 |
| Chelerythrine | 86 | 33 | 86 | 30 |
| cpd **1** | 90 | 30 | 92 | 2 |
| Pyrvinium | 86 | 44 | 69 | 1 |

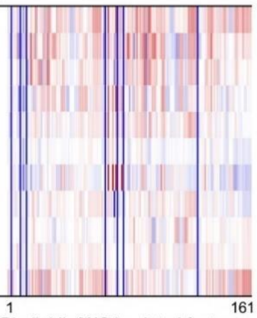


| Compound | BioSim [%] | Ind [%] | Cell count [%] | Conc [µM] |
|---|---|---|---|---|
| CPX | – | 59 | 75 | 30 |
| ML288 | 92 | 66 | 71 | 3 |
| ML288 | 83 | 54 | 65 | 10 |
| SC144 | 85 | 52 | 69 | 2 |
| GSK-J4 | 84 | 46 | 80 | 10 |
| GSK-J4 | 90 | 41 | 85 | 2 |
| JIB-04 | 70 | 45 | 79 | 2 |
| Sanguinarine | 71 | 36 | 71 | 1 |
| Chelerythrine | 91 | 33 | 86 | 30 |
| cpd **1** | 92 | 30 | 92 | 2 |
| Pyrvinium | 77 | 44 | 69 | 1 |

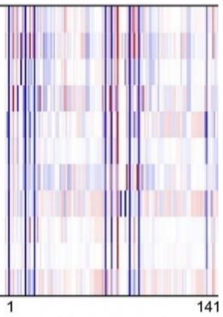


**Figure S2 (related to Figure 2): Profile comparison for ciclopirox (CPX) and biosimilar compounds** considering the features related to each stain only.

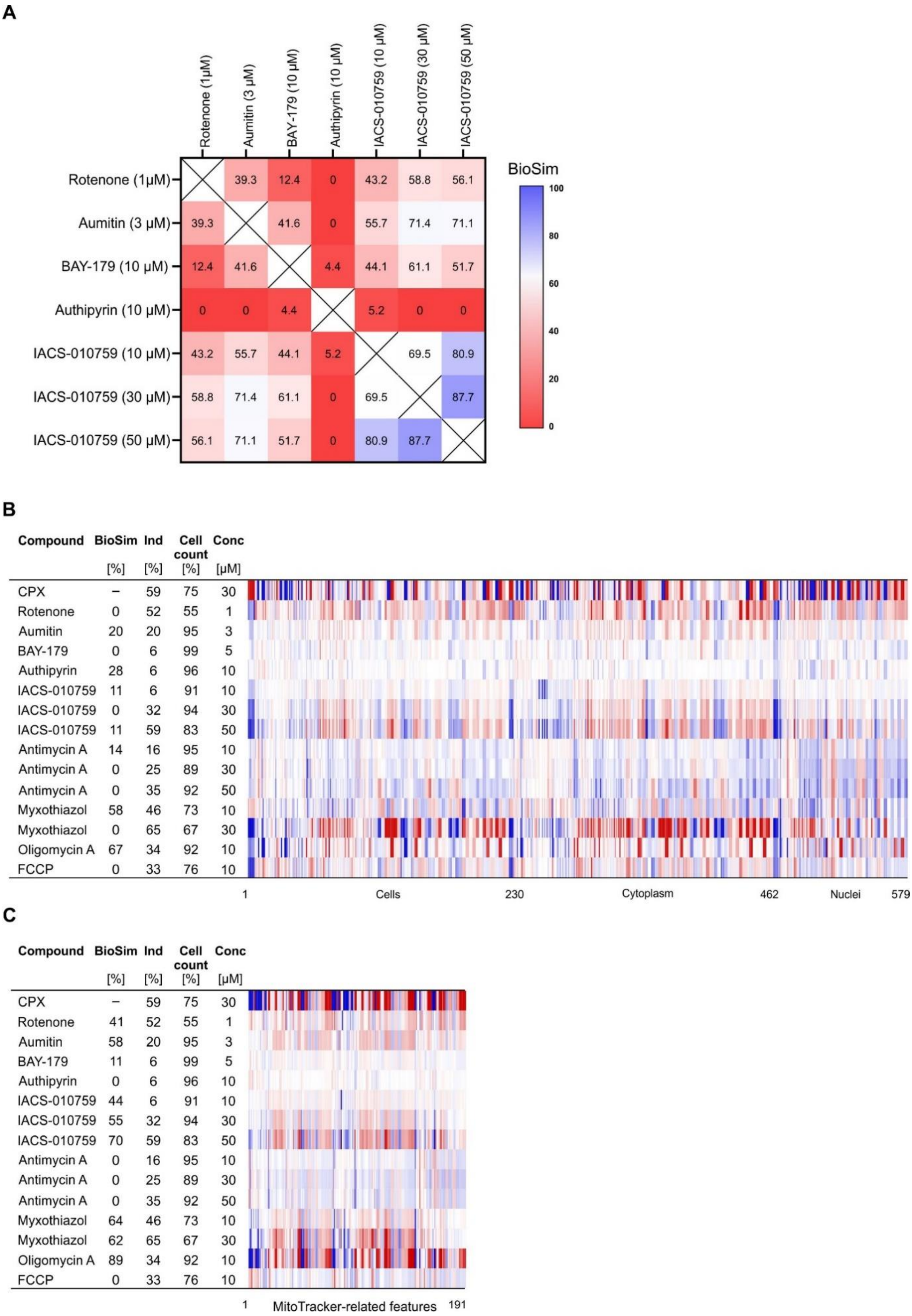


**Figure S3 (related to Figure 2): Profile analysis for inhibitors of the mitochondrial electron transport chain (ETC).** (A) Profile cross-similarity for ETC inhibitors. (B)

Comparison of the profile of ciclopirox (CPX) at 30 µM to the profiles of ETC inhibitors. (C) Comparison of the profiles for ciclopirox (CPX) at 30 µM to the profiles of ETC inhibitors considering only MitoTracker-related features. For B and C: the top line profile is set as a reference profile (100 % biological similarity, BioSim) to which the following profiles are compared. Blue color: decreased feature; red color: increased feature. Cpd: compound; BioSim: biosimilarity; Ind: induction; Conc: concentration.

**A**

| Cpd | Conc [µM] | AKT/PI3K/MTOR | Aurora | BET | DNA synthesis | HDAC | HSP90 | L/CH | $Na^+/K^+$ ATPase | Protein synthesis | PYR synthesis | Tubulin | Uncoupling |
|---|---|---|---|---|---|---|---|---|---|---|---|---|---|
| | | Cluster biosimilarities | | | | | | | | | | | |
| SB525334 | 0.2 | 0 | 0 | 51 | 0 | 54 | 0 | 15 | 65 | 28 | 0 | 46 | 16 |
| | 2 | 0 | 5 | 73 | 0 | 74 | 0 | 46 | 64 | 52 | 0 | 68 | 28 |
| | 10 | 0 | 0 | 31 | 0 | 31 | 24 | 42 | 28 | 26 | 0 | 39 | 9 |
| | 30 | 0 | 0 | 33 | 0 | 32 | 27 | 44 | 23 | 29 | 0 | 41 | 2 |
| | 50 | 0 | 0 | 30 | 0 | 31 | 33 | 41 | 19 | 32 | 0 | 39 | 2 |

**B**

| Cpd | Conc [µM] | AKT/PI3K/MTOR | Aurora | BET | DNA synthesis | HDAC | HSP90 | L/CH | $Na^+/K^+$ ATPase | Protein synthesis | PYR synthesis | Tubulin | Uncoupling |
|---|---|---|---|---|---|---|---|---|---|---|---|---|---|
| | | Cluster biosimilarities | | | | | | | | | | | |
| cpd **2** | 10 | 11 | 0 | 0 | 34 | 0 | 31 | 21 | 0 | 8 | 13 | 16 | 0 |

**C**

| Compound | BioSim [%] | Ind [%] | Cell count [%] | Conc [µM] |
|---|---|---|---|---|
| CPX | – | 59 | 75 | 30 |
| cpd **2** | 79 | 34 | 95 | 10 |

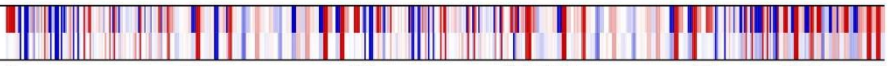


**Figure S4 (related to Figure 2): CPA profile for SB525334 and compound 2.** (A and B) Cluster biosimilarity heatmap for SB525334 (A) and compound **2** (B) Values are biosimilarity in %. (C) Comparison of the profile of ciclopirox (CPX) at 30 µM to the profile of compound **2** at 10 µM. The top line profile is set as a reference profile (100 % biological similarity, BioSim) to which the following profiles are compared. Blue color: decreased feature; red color: increased feature. Cpd: compound; BioSim: biosimilarity; Ind: induction; Conc: concentration. L/CH: Lysosomotropism/cholesterol homeostasis; PYR: pyrimidine.

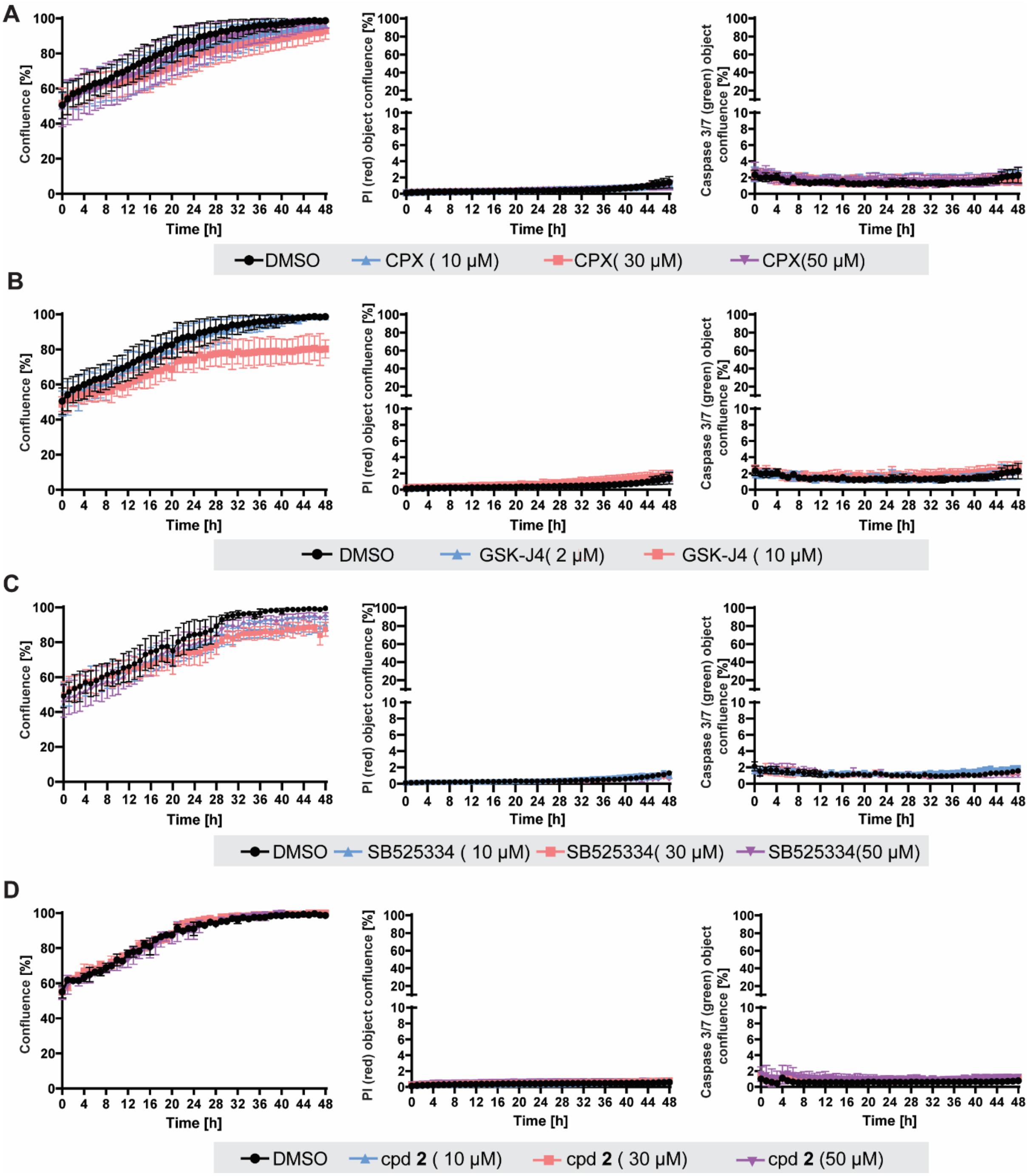


**Figure S5: Influence of the compounds on cell growth.** U-2OS cells were treated with the compounds for 48 h in presence of propidium iodide (PI) and CellEvent™ Caspase-3/7 Green to detect cell toxicity and apoptosis, respectively. Images were acquired every hour using the IncuCyte Zoom imaging system. Image-based analysis was performed to quantify cell growth by means of cell confluence as a readout, or cell toxicity and apoptosis by means of PI and caspase 3/7 activity-related fluorescence. (n =3; mean values ± SD).

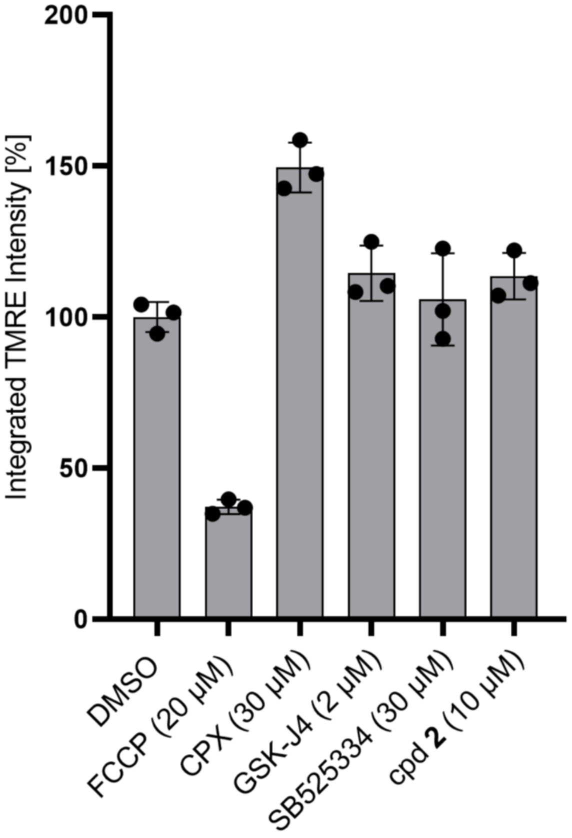


**Figure S6: Influence on the mitochondrial membrane potential.** U-2OS cells were treated with the compounds for 24 h prior to the addition of tetramethylrhodamine, methyl ester (TMRE) to determine mitochondrial membrane potential. FCCP was used as a positive control (n =3; mean values ± SD).

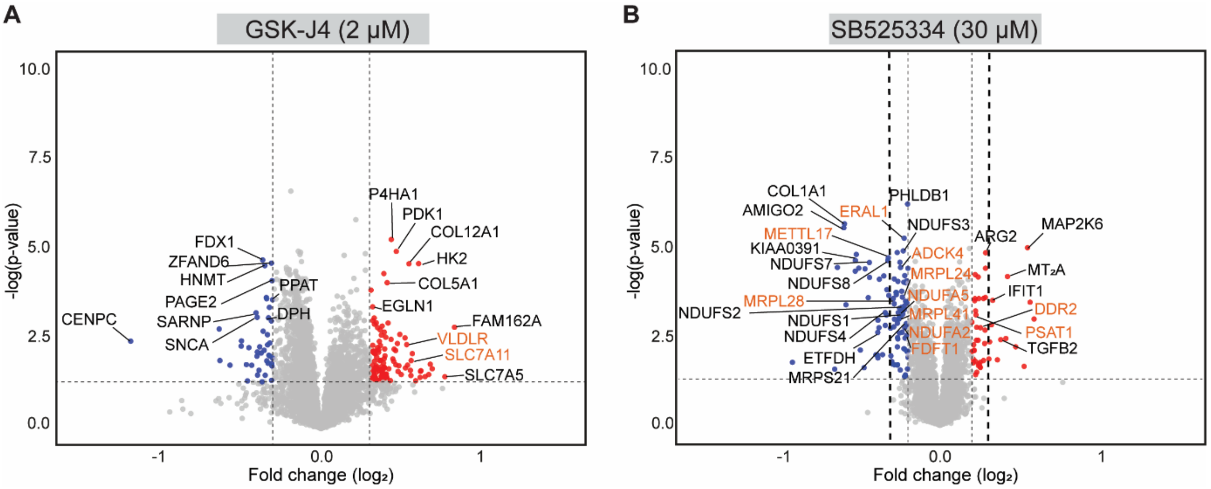


**Figure S7 (related to figure 5): Proteomics analysis.** Volcano plot of log2 fold changes in protein abundance upon treatment for 24 h with 2 µM GSK-J4 (A) or 30 µM SB525334 (B) (FC <±0.2; light gray; FC <±0.3; black). Red circles: upregulated proteins; blue circles: downregulated proteins. orange circles: proteins found regulated in Quiros et al. Volcano plots were visualized using VolcaNoseR (Goedhart and Luijsterburg, 2020). FC: fold change.

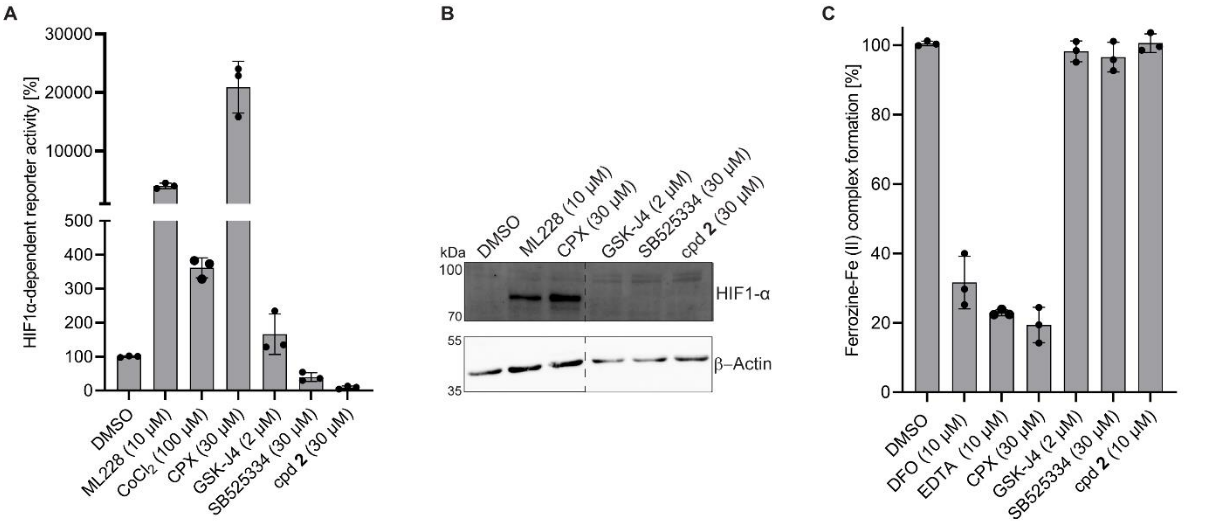


**Figure S8 (related to Figure 5): Influence of the compounds on HIF1 signaling and iron chelation.** (A) HIF1-α-dependent reporter gene assay. HEK293 cells transfected with pGL4.22-PGK1-HRE::dLUC and *Renilla* luciferase-expressing plasmids were treated with the compounds or DMSO as a control for 24 h prior to detection of firefly and *Renilla* luciferase activities. ML228 and $CoCl_2$ were used as controls for HIF1 induction. Mean values ± SD (n =3). (B) Detection of HIF1-α protein levels in U-2OS cells after treatment with the compounds for 24 h. Cells were treated with the compounds prior to detection of HIF1-α and β-actin as a reference protein using immunoblotting. Representative blot is shown (n =3). Lanes in the immunoblot were rearranged to fit the figure. See Figure S9 for the full blots. (C) Using Ferrozine-Fe(II) complex formation to determine iron chelation by the compounds. Mean values ± SD (n =3).

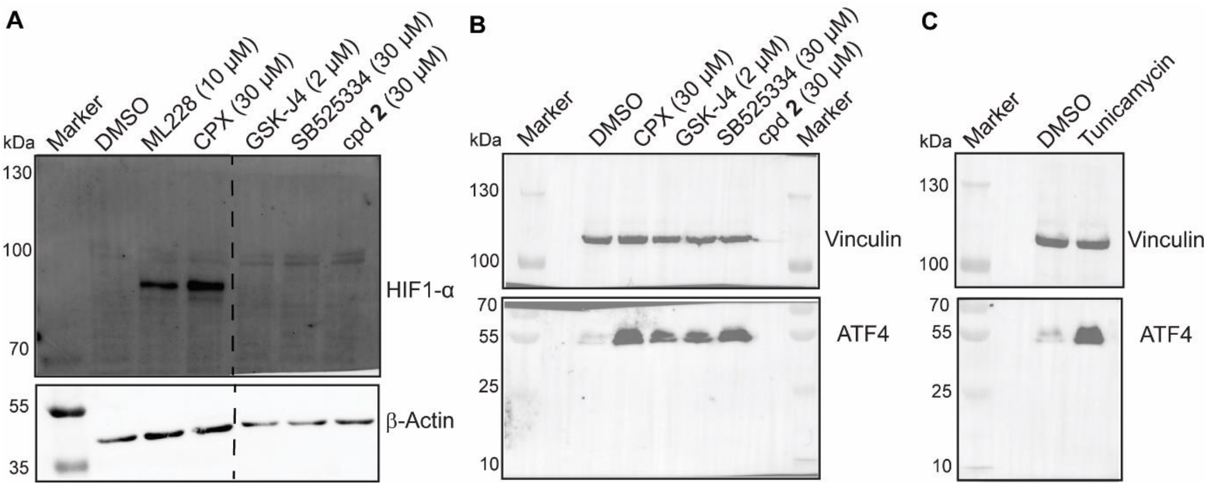


**Figure S9: Full blots for the results shown in Figure. Figure S8B (A), 6D (B) and 6E (C).**

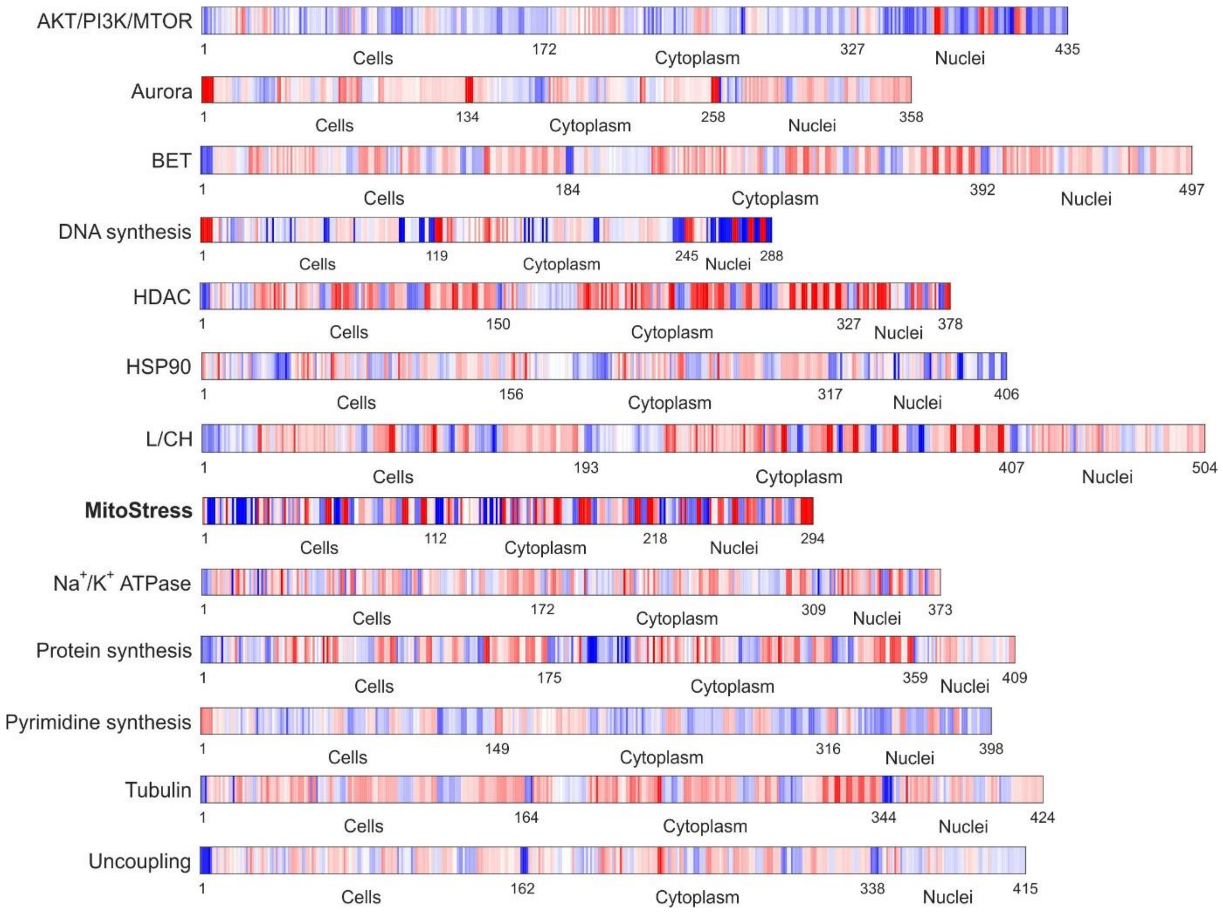


**Figure S10 (related to Figure 7): Median cluster subprofiles of the 13 defined clusters.** The median cluster subprofiles for all clusters besides MitoStress were previously reported (Pahl et al., 2023). Blue color: decreased feature, red color: increased feature.

**A**

| Cpd | Conc [µM] | AKT/PI3K/MTOR | Aurora | BET | DNA synthesis | HDAC | HSP90 | L/CH | MitoStress | Na+/K+ ATPase | Protein synthesis | PYR synthesis | Tubulin | Uncoupling |
|---|---|---|---|---|---|---|---|---|---|---|---|---|---|---|
| DFO | 3 | 38 | 19 | 0 | 92 | 0 | 0 | 0 | 43 | 0 | 0 | 81 | 0 | 0 |
| | 6 | 49 | 0 | 0 | 94 | 0 | 0 | 0 | 39 | 0 | 0 | 71 | 0 | 0 |
| | 10 | 51 | 4 | 0 | 93 | 0 | 0 | 0 | 35 | 0 | 0 | 72 | 0 | 0 |
| | 30 | 44 | 23 | 0 | 93 | 0 | 0 | 0 | 20 | 0 | 0 | 73 | 0 | 0 |
| | 50 | 46 | 13 | 0 | 94 | 0 | 0 | 0 | 30 | 0 | 0 | 65 | 0 | 0 |

Cluster biosimilarities

**B**

| Cpd | Conc [µM] | AKT/PI3K/MTOR | Aurora | BET | DNA synthesis | HDAC | HSP90 | L/CH | MitoStress | Na+/K+ ATPase | Protein synthesis | PYR synthesis | Tubulin | Uncoupling |
|---|---|---|---|---|---|---|---|---|---|---|---|---|---|---|
| Phen | 10 | 46 | 12 | 0 | 96 | 0 | 0 | 0 | 31 | 0 | 0 | 61 | 0 | 0 |
| | 30 | 44 | 17 | 0 | 96 | 0 | 0 | 0 | 63 | 0 | 0 | 60 | 0 | 0 |
| | 50 | 48 | 8 | 0 | 93 | 0 | 0 | 0 | 62 | 0 | 0 | 61 | 0 | 0 |

Cluster biosimilarities

**C**

| Cpd | Conc [µM] | AKT/PI3K/MTOR | Aurora | BET | DNA synthesis | HDAC | HSP90 | L/CH | MitoStress | Na+/K+ ATPase | Protein synthesis | PYR synthesis | Tubulin | Uncoupling |
|---|---|---|---|---|---|---|---|---|---|---|---|---|---|---|
| Deferasirox | 10 | 17 | 36 | 0 | 83 | 0 | 0 | 0 | 52 | 0 | 0 | 63 | 0 | 0 |
| | 30 | 52 | 17 | 0 | 96 | 0 | 0 | 0 | 28 | 0 | 0 | 73 | 0 | 0 |
| | 50 | 46 | 26 | 0 | 95 | 0 | 0 | 0 | 25 | 0 | 0 | 74 | 0 | 0 |

Cluster biosimilarities

**D**

| Cpd | Conc [µM] | AKT/PI3K/MTOR | Aurora | BET | DNA synthesis | HDAC | HSP90 | L/CH | MitoStress | Na+/K+ ATPase | Protein synthesis | PYR synthesis | Tubulin | Uncoupling |
|---|---|---|---|---|---|---|---|---|---|---|---|---|---|---|
| PAC-1 | 2 | 15 | 22 | 0 | 70 | 0 | 23 | 1 | 63 | 0 | 0 | 41 | 2 | 0 |
| | 6 | 38 | 30 | 0 | 92 | 0 | 0 | 0 | 41 | 0 | 0 | 75 | 0 | 0 |
| | 10 | 44 | 26 | 0 | 94 | 0 | 0 | 0 | 32 | 0 | 0 | 80 | 0 | 0 |
| | 30 | 41 | 17 | 0 | 96 | 0 | 0 | 0 | 48 | 0 | 0 | 65 | 0 | 0 |

Cluster biosimilarities

**E**

| Cpd | Conc [µM] | AKT/PI3K/MTOR | Aurora | BET | DNA synthesis | HDAC | HSP90 | L/CH | MitoStress | Na+/K+ ATPase | Protein synthesis | PYR synthesis | Tubulin | Uncoupling |
|---|---|---|---|---|---|---|---|---|---|---|---|---|---|---|
| SC144 | 2 | 41 | 9 | 0 | 93 | 0 | 16 | 0 | 91 | 0 | 0 | 59 | 0 | 0 |
| GSK-J4 | 2 | 18 | 1 | 0 | 49 | 0 | 28 | 20 | 98 | 0 | 7 | 20 | 12 | 0 |
| GSK-J4 | 10 | 15 | 0 | 2 | 44 | 0 | 26 | 27 | 98 | 0 | 11 | 9 | 22 | 0 |
| JIB-04 | 2 | 46 | 4 | 0 | 78 | 0 | 24 | 0 | 86 | 0 | 0 | 54 | 0 | 0 |
| Sanguinarine | 1 | 11 | 0 | 7 | 26 | 4 | 28 | 27 | 97 | 6 | 25 | 0 | 21 | 0 |
| Chelerythrine | 30 | 6 | 0 | 8 | 15 | 4 | 27 | 29 | 96 | 5 | 13 | 0 | 23 | 0 |
| Chelerythrine | 50 | 0 | 3 | 8 | 15 | 2 | 29 | 26 | 94 | 3 | 14 | 0 | 22 | 0 |
| ALK5i **1** | 0.2 | 0 | 0 | 11 | 0 | 7 | 30 | 26 | 81 | 10 | 17 | 0 | 25 | 0 |
| ALK5i **1** | 2 | 3 | 1 | 9 | 12 | 4 | 28 | 29 | 96 | 6 | 13 | 0 | 24 | 0 |
| Pyrivinium | 1 | 4 | 10 | 0 | 55 | 0 | 16 | 16 | 92 | 8 | 12 | 14 | 15 | 0 |

Cluster biosimilarities

**F**

| Cpd | Conc [µM] | AKT/PI3K/MTOR | Aurora | BET | DNA synthesis | HDAC | HSP90 | L/CH | MitoStress | Na+/K+ ATPase | Protein synthesis | PYR synthesis | Tubulin | Uncoupling |
|---|---|---|---|---|---|---|---|---|---|---|---|---|---|---|
| Sal003 | 3 | 17 | 0 | 23 | 0 | 26 | 56 | 17 | 10 | 11 | 30 | 0 | 30 | 31 |
| | 6 | 33 | 0 | 28 | 0 | 18 | 54 | 39 | 61 | 1 | 19 | 0 | 44 | 44 |
| | 10 | 4 | 0 | 33 | 0 | 31 | 36 | 48 | 93 | 20 | 35 | 0 | 47 | 19 |

Cluster biosimilarities

**G**

| Cpd | Conc [µM] | AKT/PI3K/MTOR | Aurora | BET | DNA synthesis | HDAC | HSP90 | L/CH | MitoStress | Na+/K+ ATPase | rotein synthesis | PYR synthesis | Tubulin | Uncoupling |
|---|---|---|---|---|---|---|---|---|---|---|---|---|---|---|
| KPL | 10 | 54 | 25 | 0 | 66 | 0 | 45 | 0 | 24 | 0 | 0 | 43 | 0 | 0 |
| | 30 | 6 | 15 | 13 | 28 | 16 | 37 | 39 | 90 | 16 | 31 | 0 | 36 | 0 |

Cluster biosimilarities

**H**

| Cpd | BioSim [%] | Ind [%] | Cell count [%] | Conc [µM] |
|---|---|---|---|---|
| Sal003 | – | 6 | 88 | 3 |
| | 49 | 23 | 83 | 6 |
| | 26 | 63 | 89 | 10 |

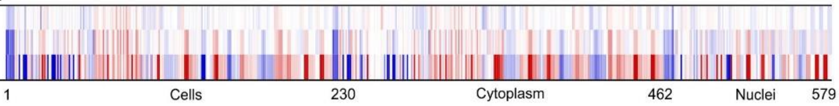


**I**

| Cpd | BioSim [%] | Ind [%] | Cell count [%] | Conc [µM] |
|---|---|---|---|---|
| KPL | – | 20 | 83 | 10 |
| | 24 | 57 | 56 | 30 |

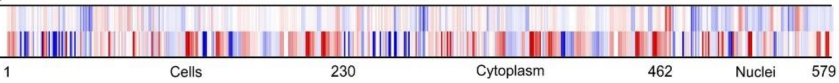


**Figure S11 (related to Figure 7): Cluster biosimilarities for DFO, phenanthroline, deferasirox, PAC-1, Sal003 and kenpaullone.** (A-F) Cluster biosimilarity heatmap for the profiles of DFO (A), phenanthroline (B), deferasirox (C), PAC-1 (D), compounds biosimilar to the ciclopirox profile at 30 µM (E), Sal003 (F) and kenpaullone (KPL) (G). (H and I) Comparison of the profiles of Sal003 (H) and kenpaullone (KPL) (I). The top line profile is set as a reference profile (100 % biological similarity, BioSim) to which the following profiles are compared. Blue color: decreased feature; red color: increased feature. Cpd: compound; BioSim: biosimilarity; Ind: induction; Conc: concentration. L/CH: Lysosomotropism/cholesterol homeostasis, PYR: pyrimidine.

**A**

| Cpd | Conc [µM] | Cluster biosimilarities | | | | | | | | | | | | |
|---|---|---|---|---|---|---|---|---|---|---|---|---|---|---|
| | | AKT/PI3K/MTOR | Aurora | BET | DNA synthesis | HDAC | HSP90 | L/CH | MitoStress | $Na^+/K^+$ ATPase | Protein synthesis | PYR synthesis | Tubulin | Uncoupling |
| Rotenone | 0.3 | 0 | 0 | 70 | 0 | 65 | 11 | 61 | 2 | 59 | 54 | 0 | 89 | 54 |
| Rotenone | 1 | 0 | 2 | 77 | 0 | 76 | 29 | 61 | 7 | 61 | 57 | 0 | 94 | 53 |
| Albendazole | 0.5 | 0 | 0 | 71 | 0 | 67 | 22 | 57 | 29 | 55 | 63 | 0 | 93 | 51 |
| Albendazole | 1 | 0 | 0 | 79 | 0 | 76 | 24 | 63 | 16 | 62 | 68 | 0 | 95 | 59 |
| Mebendazole | 0.5 | 0 | 0 | 72 | 0 | 69 | 17 | 60 | 28 | 54 | 68 | 0 | 92 | 48 |
| Mebendazole | 0.6 | 0 | 10 | 72 | 0 | 72 | 27 | 70 | 10 | 55 | 56 | 0 | 90 | 49 |
| Mebendazole | 0.2 | 0 | 33 | 44 | 0 | 44 | 17 | 39 | 5 | 26 | 32 | 0 | 73 | 28 |
| Mebendazole | 1 | 0 | 35 | 57 | 0 | 65 | 38 | 46 | 26 | 53 | 48 | 0 | 83 | 26 |
| Fenbendazole | 30 | 0 | 12 | 75 | 0 | 81 | 36 | 64 | 0 | 53 | 69 | 0 | 82 | 49 |
| Colchicine | 0.03 | 0 | 30 | 19 | 0 | 44 | 58 | 12 | 0 | 33 | 33 | 0 | 38 | 13 |
| Colchicine | 0.1 | 0 | 14 | 52 | 0 | 64 | 48 | 51 | 0 | 36 | 61 | 0 | 61 | 49 |
| Nocodazole | 0.1 | 0 | 0 | 76 | 0 | 75 | 25 | 62 | 7 | 58 | 67 | 0 | 94 | 54 |
| Digoxin | 1 | 35 | 19 | 0 | 40 | 0 | 0 | 0 | 0 | 26 | 2 | 54 | 0 | 0 |
| Dihydroouabain | 1 | 0 | 32 | 0 | 0 | 0 | 0 | 0 | 0 | 22 | 0 | 31 | 0 | 0 |
| Dihydroouabain | 3 | 25 | 2 | 0 | 14 | 0 | 0 | 0 | 0 | 35 | 4 | 23 | 0 | 7 |
| Dihydroouabain | 10 | 0 | 45 | 0 | 0 | 9 | 0 | 0 | 0 | 62 | 16 | 0 | 0 | 0 |
| Dihydroouabain | 30 | 0 | 6 | 27 | 0 | 43 | 0 | 0 | 0 | 87 | 25 | 0 | 22 | 27 |
| Oubain | 10 | 0 | 0 | 72 | 0 | 77 | 0 | 42 | 0 | 88 | 55 | 0 | 74 | 46 |
| Lovastatin | 6 | 0 | 0 | 91 | 0 | 85 | 21 | 70 | 8 | 57 | 80 | 0 | 85 | 68 |
| Lovastatin | 10 | 0 | 0 | 90 | 0 | 87 | 21 | 68 | 3 | 63 | 73 | 0 | 84 | 68 |
| Mevastatin | 10 | 0 | 0 | 61 | 0 | 66 | 0 | 59 | 0 | 54 | 21 | 0 | 47 | 45 |
| Mevastatin | 30 | 0 | 0 | 91 | 0 | 87 | 22 | 83 | 8 | 60 | 69 | 0 | 87 | 75 |
| Raloxifene | 3 | 0 | 0 | 57 | 0 | 67 | 25 | 73 | 42 | 26 | 53 | 0 | 60 | 21 |
| Raloxifene | 10 | 0 | 0 | 75 | 0 | 76 | 25 | 89 | 60 | 34 | 46 | 0 | 73 | 49 |
| Prazosin | 10 | 0 | 58 | 0 | 4 | 0 | 27 | 10 | 10 | 0 | 0 | 0 | 35 | 0 |

**B**

| Cpd | Conc [µM] | Cluster biosimilarities | | | | | | | | | | | | |
|---|---|---|---|---|---|---|---|---|---|---|---|---|---|---|
| | | AKT/PI3K/MTOR | Aurora | BET | DNA synthesis | HDAC | HSP90 | L/CH | MitoStress | $Na^+/K^+$ ATPase | Protein synthesis | PYR synthesis | Tubulin | Uncoupling |
| Enclomiphene | 1 | 0 | 0 | 57 | 0 | 60 | 34 | 71 | 59 | 15 | 44 | 0 | 58 | 39 |
| Enclomiphene | 3 | 0 | 0 | 76 | 0 | 75 | 11 | 95 | 43 | 34 | 56 | 0 | 68 | 44 |
| Enclomiphene | 10 | 0 | 0 | 84 | 0 | 81 | 12 | 89 | 35 | 42 | 69 | 0 | 78 | 61 |
| Enclomiphene | 30 | 0 | 0 | 81 | 0 | 77 | 16 | 80 | 0 | 39 | 69 | 0 | 78 | 60 |
| Amiodarone | 10 | 0 | 0 | 76 | 0 | 72 | 12 | 91 | 50 | 27 | 59 | 0 | 68 | 52 |
| Amiodarone | 30 | 0 | 0 | 76 | 0 | 73 | 21 | 87 | 59 | 34 | 61 | 0 | 74 | 52 |
| Clozapine | 10 | 0 | 0 | 76 | 0 | 70 | 6 | 79 | 7 | 43 | 33 | 0 | 65 | 52 |
| Clozapine | 30 | 0 | 0 | 77 | 0 | 77 | 28 | 92 | 46 | 33 | 52 | 0 | 72 | 48 |
| AT1 | 2 | 13 | 0 | 65 | 0 | 56 | 11 | 81 | 53 | 20 | 28 | 0 | 61 | 54 |

**Figure S12 (related to Figure 7):** Cluster biosimilarities for the profiles of compounds studied for mitotoxicity. (A) Compounds from Seal *et al*. (Seal et al., 2022) (B) Compounds from Trapotsi *et al*.(Trapotsi et al., 2022) Percent values are given. Cpd: compound; Conc: concentration; PYR: pyrimidine.

**A**

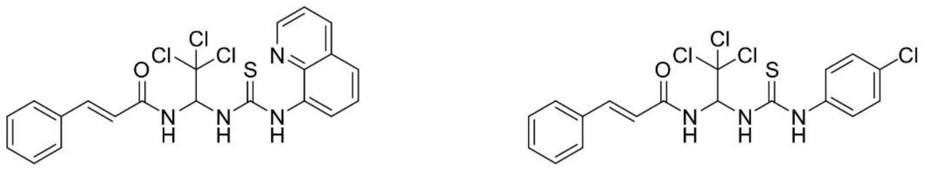

Salubrinal　　　　　　　　Sal003

**B**

| Compound | BioSim [%] | Ind [%] | Cell count [%] | Conc [μM] |
|---|---|---|---|---|
| Salubrinal | – | 7 | 98 | 10 |
| | 68 | 15 | 97 | 30 |
| | 36 | 22 | 84 | 50 |

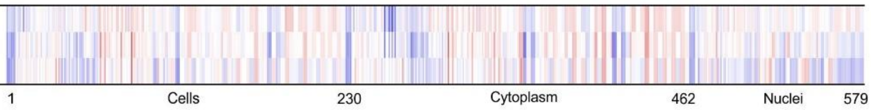


**C**

| Compound | BioSim [%] | Ind [%] | Cell count [%] | Conc [μM] |
|---|---|---|---|---|
| Sal003 | – | 6 | 88 | 3 |
| | 49 | 23 | 83 | 6 |
| | 26 | 63 | 89 | 10 |

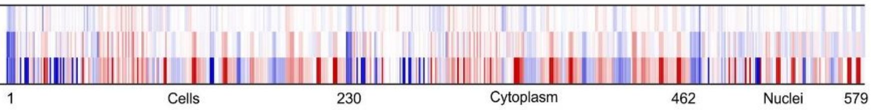


**D**

| Cpd | Conc [μM] | AKT/PI3K/MTOR | Aurora | BET | DNA synthesis | HDAC | HSP90 | L/CH | MitoStress | $Na^+/K^+$ ATPase | Protein synthesis | PYR synthesis | Tubulin | Uncoupling |
|---|---|---|---|---|---|---|---|---|---|---|---|---|---|---|
| | | Cluster biosimilarities | | | | | | | | | | | | |
| Salubrinal | 10 | 0 | 0 | 58 | 0 | 61 | 36 | 60 | 28 | 26 | 55 | 0 | 64 | 53 |
| | 30 | 0 | 0 | 66 | 0 | 71 | 49 | 56 | 0 | 41 | 54 | 0 | 63 | 71 |
| | 50 | 25 | 0 | 38 | 0 | 55 | 41 | 34 | 0 | 33 | 29 | 0 | 40 | 64 |

**E**

| | | Salubrinal 10 μM | Salubrinal 30 μM | Salubrinal 50 μM | Sal003 3 μM | Sal003 6 μM |
|---|---|---|---|---|---|---|
| Salubrinal | 30 μM | 68 | | | | |
| | 50 μM | 36 | 77 | | | |
| Sal003 | 3 μM | 40 | 51 | 48 | | |
| | 6 μM | 53 | 31 | 21 | 49 | |
| | 10 μM | 46 | 20 | 0 | 26 | 62 |

**Figure S13 (related to Figure 7): Profile analysis for salubrinal.** (A) Chemical structures of salubrinal and Sal003. (B, C) Profile similarity for salubrinal and Sal003, respectively. The top line of the heatmap profile is set as a reference profile (100 % biological similarity) to which the following profiles are compared. Blue color, decreased feature; red color, increased feature. (D) Cluster biosimilarity heatmap for salubrinal. Percent values are given. (E) Profile cross-similarity for salubrinal and Sal003. Values are biosimilarity in %. Cpd: compound; BioSim: biosimilarity; Ind: induction; Conc: concentration. L/CH: Lysosmotropism/cholesterol homeostasis; PYR: pyrimidine.

#### Supporting Tables

##### Table S2 (related to Figure 2): Compounds that are biosimilar in CPA to ciclopirox at 30 µM at the indicated concentrations.

| Name | Conc [µM] | Induction [%] | BioSim [%] | Known activity |
|---|---|---|---|---|
| Ciclopirox (olamine)<br>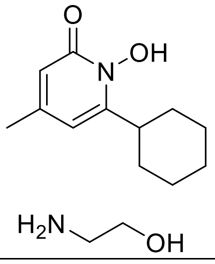 | 30 | 59.2 | 100 | Metal ion chelator |
| | 50 | 64.8 | 97.5 | |
| ML228<br>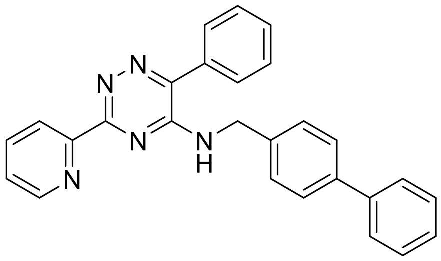 | 3 | 65.5 | 95.7 | Hypoxia Inducible Factor (HIF) pathway activator |
| | 10 | 54.4 | 92.7 | |
| NSC319726<br>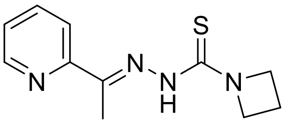 | 2 | 53.2 | 94.6 | p53(R175) mutant reactivator |
| SC144<br>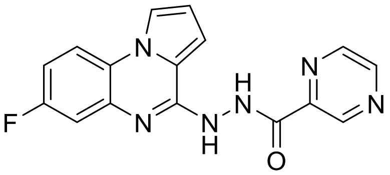 | 2 | 52.3 | 92.2 | gp130 (IL6-beta) inhibitor (Lu et al., 2020) |
| GSK-J4 (hydrochloride)<br>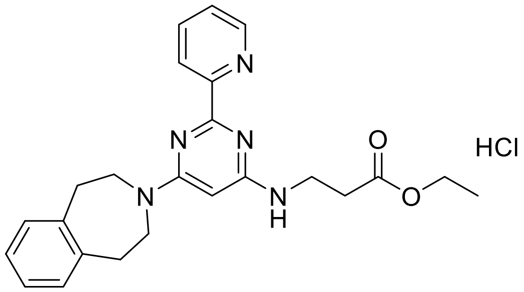 | 10 | 46.3 | 86.5 | Dual inhibitor of H3K27me3/me2-demethylases JMJD3/KDM6B and UTX/KDM6A |
| | 2 | 40.6 | 83.5 | |
| BLU9931<br>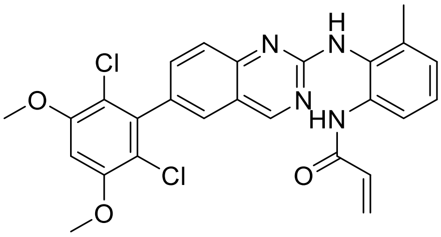 | 10 | 51.5 | 84.2 | Fibroblast growth factor receptor 4 (FGFR4) inhibitor |

| Name | Conc [µM] | Induction [%] | BioSim [%] | Known activity |
|---|---|---|---|---|
| Dephostatin 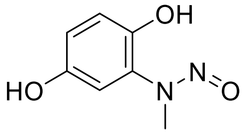 | 30 | 37 | 81.4 | CD45 protein tyrosine kinase inhibitor. |
| JIB-04 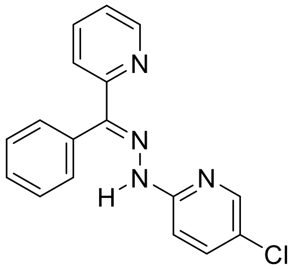 | 2 | 44.9 | 81.1 | Pan-selective Jumonji histone demethylase inihibitor |
| Sanguinarine chloride 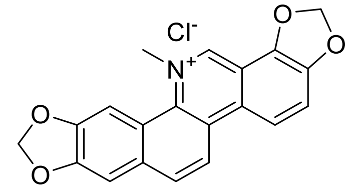 | 1 | 36.3 | 78.6 | Inhibitor of $Mg^{2+}$ and $Na^{+}$ /$K^{+}$ ATPase (Croaker et al., 2016) |
| Chelerythrine (chloride) 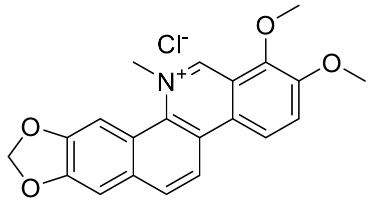 | 30 | 33.2 | 78.1 | Inhibitor of protein kinase C (Chen et al., 2022) |
| | 50 | 31.4 | 77.3 | |
| ALK5i **1** 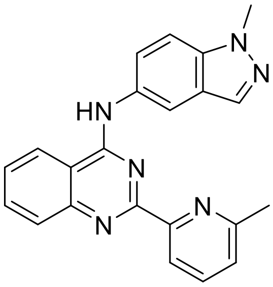 | 2 | 30.4 | 77.8 | Inhibitor of TGF-beta receptor type I (ALK5) (Gellibert et al., 2009) |
| Pyrvinium pamoate 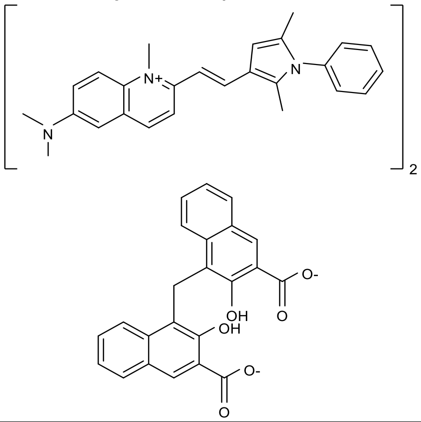 | 1 | 44.4 | 76.9 | Metabolic inhibitor (Ishii et al., 2012) |

**Table S3 (related to Figure 2). Compounds that are biosimilar in CPA to ciclopirox at 30 µM at the indicated concentrations when only MitoTracker-related features are considered**. Conc: concentration; BioSim: biosimilarity

| Trivial Name | Conc. [µM] | Induction [%] | BioSim [%] | Known activity |
|---|---|---|---|---|
| Ciclopirox (olamine) | 30.0 | 59.2 | 100.0 | metal ion chelator |
| Ciclopirox (olamine) | 50.0 | 64.8 | 99.4 | metal ion chelator |
| GSK-J4 (hydrochloride) | 10.0 | 46.3 | 97.3 | dual inhibitor of H3K27me3/me2-demethylases JMJD3/KDM6B and UTX/KDM6A |
| ML228 | 3.0 | 65.5 | 97.0 | Hypoxia Inducible Factor (HIF) pathway activator |
| Chelerythrine (chloride) | 30.0 | 33.2 | 96.7 | Inhibitor of protein kinase C |
| NSC319726 | 2.0 | 53.2 | 95.9 | p53(R175) mutant reactivator |
| SB 525334 | 50.0 | 45.9 | 95.8 | Transforming growth factor b1 receptor (ALK5) inhibitor |
| Sal003 | 10.0 | 62.9 | 95.6 | Inhibitor of the eukaryotic translation initiation factor 2a (eIF2a) phosphatase |
| Chelerythrine (chloride) | 50.0 | 31.4 | 95.5 | Inhibitor of protein kinase C |
| ML228 | 10.0 | 54.4 | 95.2 | Hypoxia Inducible Factor (HIF) pathway activator |
| Berberine (chloride hydrate) | 30.0 | 25.4 | 94.6 | Induces reactive oxygen species (ROS) generation and inhibits DNA topoisomerase |
| Berberine (chloride hydrate) | 50.0 | 31.8 | 94.5 | Induces reactive oxygen species (ROS) generation and inhibits DNA topoisomerase |
| Dynarrestin | 30.0 | 42.8 | 94.2 | Inhibitor of cytoplasmic dyneins 1 and 2 |
| SB 525334 | 30.0 | 44.7 | 93.4 | Transforming growth factor b1 receptor (ALK5) inhibitor |
| SC144 | 2.0 | 52.3 | 93.2 | gp130 (IL6-beta) inhibitor |
| Calcimycin, A23187, Calcium ionophore A23187 | 10.0 | 60.3 | 92.9 | $Ca^{2}$+ ionophore |
| Dequalinium dichloride | 10.0 | 43.0 | 92.3 | Blocker of apamin-sensitive K+ channels |
| Quinidine sulfate | 10.0 | 35.8 | 89.2 | $Na^+$ channel blocker |
| Oligomycin A | 10.0 | 34.0 | 89.2 | Mitochondrial $F_0F_1$-ATPase inhibitor |
| BLU9931 | 10.0 | 51.5 | 89.0 | Fibroblast growth factor receptor 4 (FGFR4) inhibitor |
| Kenpaullone | 30.0 | 56.8 | 88.7 | Inhibitor of CDK1/cyclin B and GSK-3b |
| Chelerythrine (chloride) | 2.0 | 17.8 | 88.5 | Inhibitor of protein kinase C |
| JIB-04 | 2.0 | 44.9 | 87.1 | Pan-selective Jumonji histone demethylase inihibitor |
| Calcimycin, A23187, Calcium ionophore A23187 | 0.3 | 61.1 | 87.1 | $Ca^{2+}$ ionophore |
| Ammonium pyrrolidinedithiocarbamate, APDC | 10.0 | 26.4 | 86.9 | Prevents induction of nitric oxide synthase (NOS) by inhibiting translation of NOS mRNA. |
| Phenserine | 30.0 | 63.6 | 86.7 | Non-competitive acetylcholinesterase (AChE) inhibitor. |
| NSC 228155 | 2.0 | 24.4 | 85.8 | Activator of EGFR |
| Halofantrine hydrochloride | 3.0 | 23.8 | 85.4 | Blocker of delayed rectifier potassium current via the inhibition of human-ether-a-go-go-related gene (HERG) channel |

**Table S7 (related to Figure 5 and 6): Overlap of downregulated proteins in Quiros et al. (Quirós et al., 2017) and this study.** The table lists proteins, which were reported by Quiros et al. to be downregulated upon the treatment with mitochondrial stressor, that were also downregulated by ciclopirox (CPX), GSK-J4, SB525334 or compound **2** (cpd **2**). Data for SB525334 is shown for log fold change (FC) of 0.3. and 0.2 as only little changes were detected at log FC of 0.2.

| 30 µM CPX | GSK-J4 | SB525334 log FC 0.3 | SB525334 log FC 0.2 | Cpd 2 |
|---|---|---|---|---|
| ADCK4 | | METTL17 | ADCK4 | DAP3 |
| ATF7IP | | | FDFT1 | FADS2 |
| CYP51A1 | | | METTL17 | FDFT1 |
| DAB2 | | | MRPL24 | MRPS12 |
| DAP3 | | | MRPL28 | MRPS7 |
| ERAL1 | | | MRPL41 | |
| FADS2 | | | NDUFA2 | |
| FDFT1 | | | NDUFA5 | |
| ICT1 | | | RAL1 | |
| MRPL11 | | | | |
| MRPL17 | | | | |
| MRPL18 | | | | |
| MRPL19 | | | | |
| MRPL24 | | | | |
| MRPL28 | | | | |
| MRPL3 | | | | |
| MRPL30 | | | | |
| MRPL40 | | | | |
| MRPL42 | | | | |
| MRPL43 | | | | |
| MRPL45 | | | | |
| MRPL51 | | | | |
| MRPS15 | | | | |
| MRPS18A | | | | |
| MRPS18B | | | | |
| MRPS2 | | | | |
| MRPS22 | | | | |
| MRPS23 | | | | |
| MRPS28 | | | | |
| MRPS35 | | | | |
| MRPS5 | | | | |
| MRPS7 | | | | |
| MRPS9 | | | | |
| MTG1 | | | | |
| NDUFA2 | | | | |
| NDUFA5 | | | | |
| NDUFA6 | | | | |
| TMEM50A | | | | |

**Table S8 (related to Figure 5 and 6): Upregulated genes and proteins in Quiros et al. (Quirós et al., 2017) and this study.** The table lists genes and proteins, which were reported by Quiros et al. to be downregulated upon the treatment with mitochondrial stressor, that were also upregulated by ciclopirox (CPX), GSK-J4, SB525334 or compound **2** (cpd **2**). Data for SB525334 is shown for log fold change (FC) of 0.3. and 0.2 as only little changes were detected at log FC of 0.2.

| 30 µM CPX | GSK-J4 | SB525334 log FC 0.3 | SB525334 log FC 0.2 | Cpd 2 |
|---|---|---|---|---|
| DNASE2<br>GRB10<br>SARS<br>VLDLR | SLC7A11<br>VLDLR | | DDR2<br>PSAT1 | AMIGO2<br>ASNS<br>CTH<br>DDR2<br>PCK2<br>PSAT1<br>PSPH<br>SESN2<br>SLC1A4<br>SLC1A5<br>SLC7A11<br>VLDLR |

#### Table S9 (related to Figure 7): MitoStress cluster defining compounds.

| Trivial_Name | Induction [%] | Conc [µM] | Cell count [%] | MitoStress cluster similarity |
|---|---|---|---|---|
| Ciclopirox (olamine) | 59 | 30 | 75 | 93 |
| Ciclopirox (olamine) | 65 | 50 | 72 | 96 |
| ML228 | 66 | 3 | 71 | 94 |
| ML228 | 54 | 10 | 65 | 95 |
| SC144 | 52 | 2 | 69 | 91 |
| GSK-J4 (hydrochloride) | 41 | 2 | 85 | 98 |
| GSK-J4(hydrochloride) | 46 | 10 | 80 | 98 |
| JIB-04 | 45 | 2 | 79 | 86 |
| Sanguinarine (chloride) | 36 | 1 | 71 | 96 |
| Chelerythrine (chloride) | 33 | 30 | 86 | 96 |
| Chelerythrine(chloride) | 31 | 50 | 79 | 94 |
| Comppound **1** | 30 | 2 | 92 | 96 |
| Pyrvinium pamoate | 44 | 1 | 69 | 92 |
| Dephostatin | 37 | 30 | 75 | 86 |
| BLU9931 | 52 | 10 | 70 | 92 |
| NSC319726 | 53 | 2 | 69 | 93 |

#### Supporting Movies

Supplementary Movie S1. U-2OS cells treated with 0.5 % DMSO as a control.

Supplementary Movie S2. U-2OS cells treated with 30 μM ciclopirox for 24 h.

Supplementary Movie S3. U-2OS cells treated with 2 μM GSK-J4 for 24 h.

#### Experimental Section

| Chemicals | Supplier | Product Number/CAS |
|---|---|---|
| CellEvent™ Caspase-3/7 Green | Thermo Fisher Scientific | Cat# C10427 |
| 1-Chlor-2,4-dinitrobenzol (CDNB) | Sigma Aldrich | Cat# 237329 |
| CellLight™ Mitochondria-GFP, BacMam 2.0 | Thermo Fisher Scientific | Cat# C10596 |
| DMEM medium (high glucose) | PAN Biotech | Cat# P04-03550 |
| DNase-free RNase A | Thermo Fisher Scientific | Cat# EN0531w |
| Ferrozine | Thermo Fisher Scientific | Cat# 10522194 |
| Fetal bovine serum | Gibco | Cat# 10500-084 |
| Hoechst 33342 | Cell signalling | Cat #4082S |
| HRP (Goat anti-Rabbit IgG) | Thermo Fisher Scientific | Cat# 31460 |
| IRDe 680RD (donkey anti-mouse) | Li-Cor Biosciences | Cat# 26-68072 |
| IRDye 800CW (donkey anti-rabbit) | Li-Cor Biosciences | Cat# 926-32213 |
| IRDye 800CW (goat anti-mouse) | Li-Cor Biosciences | Cat# 926-32210 |
| Iron (II) sulfate heptahydrate | Sigma Aldrich | Cat# F8633 |
| Lipofectamine 2000 | Thermo Fisher Scientific | Cat# 11668030 |
| Mito Tracker Deep Red | Thermo Fisher Scientific | Cat# M22426 |
| MitoSOX™ Red dye | Thermo Fisher Scientific | Cat# M36008 |
| Mouse monoclonal anti-HIF1-α | Novus | Cat# NB100-105 |
| Non-essential amino acids | PAN Biotech | Cat# P08-32100 |
| Propidium iodide | Sigma Aldrich | Cat# P4864 |
| Rabbit monoclonal anti ATF4 | Cell Signaling | Cat# 11815 |
| Seahorse XF Calibrate | Agilent | 100840-000 |
| Seahorse XF DMEM medium pH 7.4 | Agilent | 103575-100 |
| Sodium pyruvate | PAN Biotech | Cat# P04-43100 |
| Sso Advanced Universal SYBR Green Supermix | Bio-Rad | Cat# 1725274 |
| Tetramethylrhodamine ethyl ester (TMRE) | Thermo Fisher Scientific | Cat# T669 |
| **Commercial Kits** | **Supplier** | **Product Number** |
| Cell Mito Stress Test kit | Agilent | 103010-100 |
| DC Protein Assay Kit II: | Bio-Rad | 000112 |
| Dual-Luciferase Reporter Assay System | Promega | E1960 |
| Quanti Tect Reverse Transcription Kit | Qiagen | 205313 |
| Qubit RNA BR Assay Kit | Thermo Fisher scientific | Q10210 |
| MycoAlert™ mycoplasma detection kit | Lonza | LT07-318 |
| Seahorse XFp Mito Stress Test Kit | Agilent | 103010-100 |
| **Cell lines** | **Supplier** | |
| HEK293Tcells (humane mbryonic kidney cells) | ATCC | CRL1268 |
| Human U-2OS cells (female) | CLS | Cat# 300364; RRID:CVCL_0042 |
| **Software** | | |
| ChemDraw 22.2.0 | PerkinElmer | |

| | | |
|---|---|---|
| Fiji imageJ 1.52 | | RRID:SCR_003070; https://imagej.net/ |
| GraphPad Prism 9.0 | GraphPad | RRID:SCR_002798: https://www.graphpad.com |
| Image Lab 6.0.1 | Bio-Rad | |
| IncuCyte Zoom | Essen BioScience | https://www.essenbioscience.com/en/products/incucyte |
| Ingenuity Pathway Analysis (IPA) | Qiagen | |
| Leica Application Suite Advanced Fluorescence | Leica | |
| MetaMorph 7.7.8.0 | Molecular Devices | RRID:SCR_002368; http://www.moleculardevices. com/Products/Software/Meta-Imaging-Series/ MetaMorph.html |
| Seahorse Wave | Agilent | |
| ZEN 2011 software | Carl Zeiss | RRID: SCR_013672 https://www.zeiss.com/microscopy/en/products/software/zeiss-zen.html |
| **Devices** | | |
| Automated Screening Microscope | Zeiss | Axiovert 200M |
| Broadband Confocal Microscope | Leica | TCS SP5 |
| IncuCyte® Zoom | Essen BioScience | https://www.essenbioscience.com/en/products/incucyte |
| Sparks plate reader | Tecan | https://lifesciences.tecan.com/microplate-readers |
| Seahorse XFp analyzer | Agilent | |

#### Cell culture

The U-2OS female human bone osteosarcoma cell line was cultured in Dulbecco's Modified Eagle's medium (DMEM, high glucose) supplemented with 4°mM L-glutamine, 10°% fetal bovine serum, 1°mM sodium pyruvate and non-essential amino acids. The cells were incubated at 37°C and 5°% $CO_2$ in humidified atmosphere. The MycoAltert$^{TM}$ Mycoplasma Detection Kit was used monthly according to the manufacturer's instruction to detect contamination with mycoplasma. Cell were always tested free of mycoplasma.

#### Cell Painting assay

The Cell Painting Assays follows closely the method described by Bray et al.(Bray et al., 2016) as recently reported (Pahl et al., 2023). "Initially, 5 µl U2OS medium were added to each well of a 384-well plate (PerkinElmer CellCarrier-384 Ultra). Subsequently, U-2OS cell were seeded with a density of 1600 cells per well in 20 µL medium. The plate was incubated for 10 min at the ambient temperature, followed by an additional 4 h incubation (37°C, 5 % $CO_2$). Compound treatment was performed with the Echo 520 acoustic dispenser (Labcyte). Different concentrations of DMSO were used as controls dependent on the used compound concentration, e.g., 0.1 % DMSO was used as a control for the profiling of compounds at 10 µM. Samples at a given compound concentration were compared to the DMSO sample of the same DMSO concentration. Incubation with compound was performed for 20 h (37°C, 5 % $CO_2$). Subsequently, mitochondria were stained with Mito Tracker Deep Red (Thermo Fisher Scientific, Cat. No. M22426). The Mito Tracker Deep Red stock solution (1 mM) was diluted to a final concentration of 100 nM in prewarmed medium. The medium was removed from the plate leaving 10 µl residual volume and 25 µl of the Mito Tracker solution were added to each well. The plate was incubated for 30 min in darkness (37°C, 5 % $CO_2$). To fix the cells 7 µl of 18.5 % formaldehyde in PBS were added, resulting in a final formaldehyde concentration of 3.7 %. Subsequently, the plate was incubated for another 20 min in darkness (RT) and washed three times with 70 µl of PBS. (Biotek Washer Elx405). Cells were permeabilized by addition of 25 µl 0.1 % Triton X-100 to each well, followed by 15 min incubation (RT) in darkness. The cells were washed three times with PBS leaving a final volume of 10 µl. To each well 25 µl of a staining solution were added, which contains 1 % BSA, 5 µl/ml Phalloidin (Alexa594 conjugate, Thermo Fisher Scientific, A12381), 25 µg/ml Concanavalin A (Alexa488 conjugate, Thermo Fisher Scientific, Cat. No. C11252), 5 µg/ml Hoechst-33342 (Sigma, Cat. No. B2261-25 mg), 1.5 µg/ml WGA-Alexa594 conjugate (Thermo Fisher Scientific, Cat. No. W11262) and 1.5 µM SYTO 14 solution (Thermo Fisher Scientific, Cat. No. S7576). The plate is incubated for 30 min (RT) in darkness and washed three times with 70 µl PBS. After the final washing step, the PBS was not aspirated. The plates were sealed and centrifuged for 1 min at 500 rpm.

The plates were prepared in triplicates with shifted layouts to reduce plate effects and imaged using a Micro XL High-Content Screening System (Molecular Devices) in 5 channels (DAPI: Ex350-400/ Em410-480; FITC: Ex470-500/ Em510-540; Spectrum Gold: Ex520-545/ Em560-585; TxRed: Ex535-585/ Em600-650; Cy5: Ex605-650/ Em670-715) with 9 sites per well and 20x magnification (binning 2).

The generated images were processed with the *CellProfiler* package (https://cellprofiler.org/, version 3.0.0)(Carpenter et al., 2006) on a computing cluster of the Max Planck Society to extract 1716 cell features per microscope site. The data was then further aggregated as medians per well (9 sites -> 1 well), then over the three replicates.

Further analysis was performed with custom *Python* (https://www.python.org/) scripts using the *Pandas* (https://pandas.pydata.org/) and *Dask* (https://dask.org/) data processing libraries as well as the *Scientific Python* (https://scipy.org/) package.

From the total set of 1716 features, a subset of highly reproducible and robust features was determined using the procedure described by Woehrmann et al.(Woehrmann et al., 2013) in the following way:

Two biological repeats of one plate containing reference compounds were analysed. For every feature, its full profile over each whole plate was calculated. If the profiles from the two repeats showed a similarity >= 0.8 (see below), the feature was added to the set.

This procedure was only performed once and resulted in a set of 579 robust features out of the total of 1716 that was used for all further analyses.

The phenotypic profiles were compiled from the Z-scores of all individual cellular features, where the Z-score is a measure of how far away a data point is from a median value.

Specifically, Z-scores of test compounds were calculated relative to the Median of DMSO controls. Thus, the Z-score of a test compound defines how many MADs (Median Absolute Deviations) the measured value is away from the Median of the controls as illustrated by the following formula:

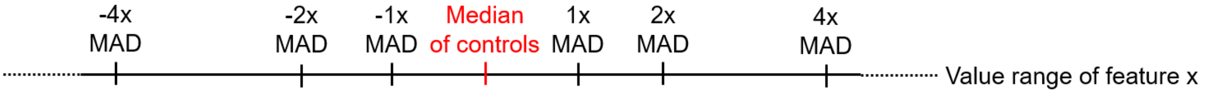

$$z - score = \frac{value_{meas.} - Median_{Controls}}{MAD_{Controls}}$$

The phenotypic compound profile is then determined as the list of Z-scores of all features for one compound.

In addition to the phenotypic profile, an induction value was determined for each compound as the fraction of significantly changed features, in percent:

$$Induction\ [\%] = \frac{number\ of\ features\ with\ abs.\ \ values\ > 3}{total\ number\ of\ features}$$

Similarities of phenotypic profiles (termed *Biosimilarity*) were calculated from the correlation distances (CD) between two profiles (https://docs.scipy.org/doc/scipy/reference/generated/scipy.spatial.distance.correlation.html) (Grigalunas et al., 2021):

$$CD = 1 - \frac{(u - \bar{u}) \cdot (v - \bar{v})}{\|(u - \bar{u})\|_2 \|(v - \bar{v})\|_2}$$

where $\bar{x}$ is the mean of the elements of $x$, $x \cdot y$ is the dot product of $x$ and $y$, and $\|x\|_2$is the Euclidean norm of $x$:

$$\|x\|_2 = \sqrt{x_1^2 + x_2^2 + \cdots + x_n^2}$$

The Biosimilarity is then defined as:

$$Biosimilarity = 1 - CD$$

Biosimilarity values smaller than 0 are set to 0 and the Biosimilarity is expressed in percent (0-100)."

**CPA subprofile analysis**

Cluster subprofiles were generated as recently described (Pahl et al., 2023). For a set of cluster defining profiles, dominating features were extracted as follows: for each profile, the sign for each of the 579 features value was assessed. The counter for positive or negative values was determined. For all cluster-defining compounds, the maximum of the two counters was determined and divided by the total number of defining profiles. A given feature was added to the cluster profile if its value has the same sign (i.e., positive or negative feature values) for 85 % of the defining profiles. Afterwards, a representative median subprofile for the cluster was calculated by taking the median values over all cluster-defining profiles for every given feature and combining them into a new reduced profile. This median (*consensus*) subprofile was then used to calculate the biosimilarity of test compounds to the defined cluster subprofiles. Due to the shorter cluster subprofiles, the cluster biosimilarity threshold was set to 80 %.

##### Mito tracker assay

U-2OS cells were seeded at a density of 5,000 cells/well into black 96-well plates with clear, flat bottom and incubated overnight at 37°C and 5 % $CO_2$. Supernatant was exchanged for test compound-containing medium, followed by 60 min of incubation at 37°C and 5 % $CO_2$. Mito Tracker Deep Red and Hoechst-33342 were added with final concentrations of 100 nM and 5 mg/ml, respectively. Cells were incubated for 3 min at 37 C and 5 % $CO_2$, rinsed twice with PBS and fixed in 4 % paraformaldehyde in PBS for 5 min at room temperature. Cells were imaged in PBS at 10x magnification using an Axiovert 200 M automated fluorescence microscope (Carl Zeiss, Germany). Stain intensities per cell were analysed using MetaMorph 7.7.8.0. Data were normalized to the value for cells that were treated with DMSO, which was set to 100 %.

##### Visualization of the mitochondrial network

One day prior to compound treatment, 3 µl of CellLight™ Mitochondria-GFP BacMam 2.0 reagent was added per $1x10^4$ U-2OS cells directly to the complete medium. 3,000 cells/well were seeded in 8 well ibidi chambers and incubated overnight at 37°C and 5°% $CO_2$. The next day, the compounds were added to the cells and the fluorescence of live cells was recorded in real-time for 24°h using a confocal microscope SP5 Leica at excitation / emission 488/555°nm for 24°h at 37°C and 5°% $CO_2$). The images and movies were analyzed using FIJI ImageJ version 1.52 (Rasband et al., 2015).

##### Real-time live-cell analysis

Cell growth and compound toxicity were observed by real-time live-cell analysis using the IncuCyte Zoom (Essen Bioscience). 5,000 U-2OS cells were seeded per well in black 96-well plate and incubated at 37°C and 5°% $CO_2$ overnight. The medium was then exchanged with fresh medium containing the compounds or DMSO as a control. Propidium iodide and CellEvent™ Caspase-3/7 Green were also added to the medium to monitor compound toxicity and apoptosis over time. Cells were incubated for 48°h and were imaged every hour. Cell confluence was quantified as a measure of cell growth using the IncuCyte® ZOOM software (2018A). Red object confluence and green object confluence was quantified as a measure of PI-positive cells and caspase-3/7 activity, respectively.

**Proteome Profiling**

$1x10^6$ U-2OS cells were seeded into a T75 flask. A day after, the medium was exchanged with DMEM containing of compound (10°µM and 30°µM ciclopirox; 2°µM GSK-J4, 30°µM SB-525334 and 30°µM compound°**2**). DMSO was used as a control. After incubation at 37°C and 5°% $CO_2$ for 24°h, the medium was removed and cells were washed with PBS. Cells were detached and washed twice with ice-cold PBS followed by centrifugation. Cells were resuspended in 1°ml PBS containing protease inhibitors and were lysed by freeze-thawing followed by centrifugation for 20°min at 15,000xg. supernatants were collected and protein concentration was determined using a DC assay. 75°µl of 2°g/l cell lysate was mixed with 75°µl 200°mM triethylammonium bicarbonate buffer (TEAB) buffer. After addition of 7.5°µl TCEP and incubation at 55°C for 30°min, samples were treated with 7.5°µl iodoacetamide (375°mM) for 30°min in the dark. By adding 900°µl prechilled acetone, proteins were precipitated and incubation at -20°C overnight. The next day, samples were centrifuged for 10°min at 8,000xg and 4°C and supernatants were removed. The dry protein pellet was dissolved in TEAB buffer containing trypsin, which was used according to manufacturer's protocol. After incubation overnight at 37°C samples were labeled with TMT label according to the manufacturer's instruction. All experiments were performed in biological triplicates.

Prior to nanoHPLC-MS/MS analysis samples were fractionated into 10 fractions on a C18 column using high pH conditions to reduce the complexity of the samples and thereby increasing the number of quantified proteins. Therefore, samples were dissolved in 120°µl of 20°mM ammonium formate ($NH_4COOH$) at pH 11, followed by ultrasonication for 2°min, subsequent vortexing for 1°min and centrifugation at 8,000xg for 3°min at room temperature. 50 µl of the C18 supernatant was injected onto a XBridge C18 column (130 Å, 3.5°µm, 1mm°x 150°mm) using a U3000 capHPLC system (ThermoFisher scientific, Germany). Separation was performed at a flow rate of 50°µl/min using 20 mM $NH_4COO$ pH 11 in water as solvent A and 40°% 20°mM $NH_4COO$ pH 11 in water premixed with 60°% acetonitrile as solvent B. Separation conditions were 95°% solvent A/5°% solvent B isocratic for the first 10°min, to desalt the samples, followed by a linear gradient up to 25°% in 5°min, a second linear gradient up to 65°% solvent B in 60°min, and a third linear gradient up to 100°% B in 10°min. Afterwards the column was washed at 100°% solvent B for 14°min and re-equilibrated to starting conditions. Detection was carried out at a valve length of 214°nm. The eluate between 15 and 100°min was fractionated into 10 fractions (30°s per fraction, circular fractionation using 10 vials). Each fraction was dried in a SpeedVac at 30°C until complete dryness and subsequently subjected to nanoHPLC-MS/MS analysis.

For nanoHPLC-MS/MS analysis, samples were dissolved in 20°µl of 0.1°% TFA in water and 1-3 µl were injected onto an UltiMateTM 3000 RSLCnano system (ThermoFisher scientific, Germany) online coupled to a Q Exactive™ HF Hybrid Quadrupole-Orbitrap Mass Spectrometer equipped with a nanospray source (Nanospray Flex Ion Source, Thermo Scientific). All solvents were LC-MS grade. To desalt the samples, they were injected onto a pre-column cartridge (5°µm, 100°Å, 300 µm°ID * 5°mm, Dionex, Germany) using 0.1°% TFA in water as eluent with a flow rate of 30°µl/min. Desalting was performed for 5 min with eluent flow to waste followed by back-flushing of the sample during the whole analysis from the pre-column to the PepMap100 RSLC C18 nano-HPLC column (2°µm, 100°Å, 75°µm ID × 50°cm, nanoViper, Dionex, Germany) using a linear gradient starting with 95°% solvent A (water containing 0.1°% formic acid)/5°% solvent B (acetonitrile containing 0.1°% formic acid) and increasing to 60°% solvent A 0.1°% formic acid / 40°% solvent B in 120°min using a flow rate of 300°nl/min. Afterwards the column was washed (95°% solvent B as highest acetonitrile concentration) and re-equilibrated to starting conditions. The nano-HPLC was online coupled to the Quadrupole-Orbitrap Mass Spectrometer using a standard coated SilicaTip (ID 20°µm, Tip-ID 10°µM, New Objective, Woburn, MA, USA).

A mass range of m/z 300 to 1650 was acquired with a resolution of 60000 for full scan, followed by up to 15 high energy collision dissociation (HCD) MS/MS scans of the most intense at least doubly charged ions using a resolution of 30000 and a NCE energy of 35°%. Data evaluation was performed using MaxQuant software (1.6.17.0) (Cox and Mann, 2008). including the Andromeda search algorithm and searching the human reference proteome of the Uniprot database. The search was performed for full enzymatic trypsin cleavages allowing two miscleavages. For protein modifications carbamidomethylation was chosen as fixed and oxidation of methionine and acetylation of the N-terminus as variable modifications. For relative quantification the type “reporter ion MS2” was chosen and for all lysines and peptide N-termini TMT labels were defined. The mass accuracy for full mass spectra was set to 20°ppm (first search) and 4.5°ppm (second search), respectively and for MS/MS spectra to 20°ppm. The false discovery rates for peptide and protein identification were set to 1°%. Only proteins for which at least two peptides were quantified were chosen for further validation. Relative quantification of proteins was carried out using the reporter ion MS2 algorithm implemented in MaxQuant. The proteinGroups.txt file was used for further analysis. All proteins which were not identified with at least two razor and unique peptides in at least one biological replicate were filtered off. The replicates were grouped together and all proteins not quantified in at least three replicates in at least one of the groups (treated or control, respectively) were filtered off. Afterwards these values were normalized to the median and a two-sides *t*-test was performed. Only proteins with a *p*°value<°0.03 and *p* value>-°0.03 were considered as statistically significantly up-or down regulated. For compound SB525334 both p°value<°0.03 and *p*°value>-°0.03 and *p* value< 0.02 and *p*°value>-°0.02 were considered. Pathway-over representation analysis was performed using the Ingenuity Pathway Analysis tool (IPA, Qiagen Version 70750971). Volcano plots were generated using the web-based tool VolcaNoseR (Goedhart and Luijsterburg, 2020).

**Seahorse Cell Mito Stress Test**

The Cell Mito Stress Test kit was performed using Seahorse XFp analyser (Agilent, USA) according to the manufacturer's protocol. $2x10^5$ U-2OS cells were seeded per wells into XFp cell culture mini plate and incubated overnight at 37°C, 5°% $CO_2$. Using the XF Calibrant, the XFp cartridges were hydrated and incubated overnight at 37°C. The next day, cell medium was exchanged to pH 7.4 DMEM-based assay medium (Agilent, USA) containing 2°mM GlutaMAX (ThermoFisher), 1°mM sodium pyruvate (PAN Biotech, Germany) and 25°mM glucose (SigmaAldrich, Germany). After five measurement of baseline recording, the test compounds were injected, followed by ten measurement intervals. Subsequently, oligomycin A, FCCP and rotenone/antimycin A were injected, followed by three measurement intervals after each injection. For compound pretreatment, the test compound was added 24°h before the assay to the XFp cell culture plates. The background was subtracted from all data and values were normalized to the last baseline measurement (which was set to 100°%) using the Wave software Version 2.6.0 (Agilent, USA). The results were plotted using GraphPad Prism 9 software.

**Analysis of the mitochondrial membrane potential**

$5°x°10^3$ U-2OS cells per well were seeded in a black 96-well plate with clear flat bottom and incubated for 24°h at 37°C and 5°% $CO_2$. Afterwards, the seeding medium was replaced with medium containing the compounds. After incubation for 24°h at 37°C and 5°% $CO_2$, 20°µM FCCP or 0.5°% DMSO were added as controls. Cells were incubated at 37°C and 5°% $CO_2$ for 10 more minutes. The staining solution was prepared by adding 1°µM TMRE and 5°µg/ml Hoechst-33342 to DMEM supplemented with 4°mM L-glutamine, 10°% fetal bovine serum, 1°mM sodium pyruvate and non-essential amino acids. After removing the medium from the cells, the staining solution was added, and the cells were incubated for 30°min at 37°C and

5°% $CO_2$. Then, the solution was removed, and the cells were washed twice with PBS, before 0.2°% BSA in PBS was added. TMRE fluorescence was recorded using the Axiovert 200M automated microscope (Carl Zeiss, Germany) with excitation/emission wavelength of 549°nm/575°nm for TMRE and 361°nm/497 nm for Hoechst-33342 at a tenfold magnification and at 37°C and 5°% $CO_2$. The obtained images were analyzed using the Multi Wavelength Cell Scoring function of the Meta Morph software version 7.7.8.0 and the results were plotted using GraphPad Prism 9 software.

**MitoSOX Red Assay**

Mitochondrial superoxide levels were determined using the indicator dye MitoSOX Red (Thermo Fisher, USA). 15,000 U-2OS were seeded per well into black 96-well plates with clear flat bottom and incubated at 37°C and 5°% $CO_2$ overnight. Seeding medium was exchanged for staining medium comprising DMEM without additives and containing 5°µM MitoSOX Red and 5°µg/µl Hoechst-33342 (ThermoFisher, USA). Cells were incubated for 30°min at 37°C and 5°% $CO_2$. Subsequently, the medium was exchanged for DMEM with additives containing the test compounds, followed by 60°min of incubation at 37°C and 5°% $CO_2$. Cells were fixed in PBS containing 0.5°% paraformaldehyde for 10°min at room temperature and washed three times with PBS. Cells were imaged using an Axiovert 200M automated microscope (Carl Zeiss, Germany) at 10x magnification. MetaMorph 7.7.8.0 (Visitron, Germany) was used to quantify the integrated fluorescence intensity of MitoSOX Red per cell. The data was normalized to control cells treated with either DMSO (set to 0°%) or 10°µM 1-Chlor-2,4-dinitrobenzol (CDNB) (set to 100°%). The results were plotted using GraphPad Prism 9 software.

**Reverse Transcriptase-Quantitative PCR (RT-qPCR)**

U-2OS cells were seeded into 6-well plates (1x$10^{5}$°cells/well) and incubated for 24°h until they reached approximately 80°% confluence. Cells were then treated with the compounds or DMSO as a control for 24°h. The total RNA was isolated using the RNAeasy Kit (Qiagen, #74104) including the DNase digestion step. The concentration of RNA was determined by using the RNA BR assay kit from (ThermoFisher, #Q10210) in conjunction with the Qubit4.0 (ThermoFisher). cDNA was obtained using the QuantiTect Reverse Transcription Kit (Qiagen, #205313). The relative mRNA amount of the ATF4 gene was evaluated using the SsoAdvanced Universal SYBR Green Supermix (Bio-Rad, #°1725274) using the CFX96 Real-Time PCR Detection System (Bio-Rad, Germany). Relative expression levels were calculated using the ΔΔCt method (Pfaffl, 2001), using *Gapdh* as a reference gene. Gene expression levels for samples that were treated with DMSO were set to 100°%. The results were plotted using GraphPad Prism 9 software. Employed primer pairs (obtained from Sigma-Aldrich) were as follows: Human-ATF4 (NM_001675) fw: 5′- TTCTCCAGCGACAAGGCTAAGG-3′, rv: 5′-CTCCAACATCCAATCTGTCCCG-3′. Human-GAPDH (NM_002046) fw: 5′-GTCTCCTCTGACTTCAACAGCG-3′, rv: 5′- ACCACCCTGTTGCTGTAGCCAA-3′.

#### Iron chelation assay

Compounds were incubated with 12.5°µM Fe(II) ($FeSO_4$) at room temperature for 10°min in a clear 96-well white plate. DMSO, deferoxamine and EDTA were used as controls. Afterwards, 0.5°mM ferrozine was added to the solution, and the absorbance at 561°nm was detected using a TecanSpark® Microplate Reader. The results were plotted using GraphPad Prism 9 software.

#### HIF1-α Reporter Gene Assay

HEK-293T cells were transfected with pGL4.22-PGK1-HRE::dLUC plasmid (Promega,#9PIE412) and pRL-TK plasmid for constitutive expression of *Renilla* luciferase (Promega,#E2241). The plasmids were added to Opti-MEM medium in a ratio of 10:1. 3 µl/µg. The Lipofectamine 2000 transfected reagent were added to the solution and incubated for 15 min and then 2.5 x $10^4$ cells/well were seeded into 96-well white plate (Corning, #353075). Then, the cells were incubated for 24 h prior to compound treatment. 100 µM $CoCl_2$ and 10 µM ML228 were used as positive controls for 24 h. Luciferase activities were determined using the Dual-Glo Luciferase Assay System (Promega, #E2940). Values obtained for the firefly luciferase were normalized to the corresponding *Renilla* luciferase values. Results are shown as fold induction determined upon normalization to the DMSO control. The results were plotted using GraphPad Prism 9 software.

#### Immunoblotting

For quantification of the HIF1-α and ATF4 protein levels, U-2OS cells were seeded into 6-well plates and incubated until they reached a confluence of 80 %. After incubation for 24 h at 37°C and 5 % $CO_2$, cells were treated with different concentrations of the compounds or DMSO as a control. After incubation for 24 h at 37°C and 5 % $CO_2$, cells were washed with PBS followed by detachment using a cell dissociation solution (Gibco, # 13151-014) for 10 min at 37°C. Detached cells were collected in 1.5 ml low-binding Eppendorf tubes (Eppendorf, #0030108116). Samples were then centrifuged for 3°min at 340xg, and the cell pellets were washed with ice-cold PBS. For HIF1-α, cells were suspended in a lysis buffer (0.01 % Bromphenol blue, 10 % Glycerol, 20 % SDS, 62.5 mM Tris (pH 6.8), and 5 % 2-mercaptoethanol). For detection of ATF4, cells were lysis in RIPA buffer. Three freeze−thaw cycles were performed to lyse the cells. Samples were centrifuged at 16,000xg and 4°C for 30°min, supernatants were transferred to fresh low-binding Eppendorf tubes, and protein concentrations were determined using the DC protein assay according to the instructions of the manufacturer. Proteins were separated via SDS-PAGE and using wet blotting proteins were transferred onto a polyvinylidene difluoride (PVDF) membrane. Membranes were stained for HIF1-α (BD Biosciences, #610959, 1:500 dilution), and β-actin as a control (Abcam, #ab8227). For quantification of ATF4 levels, membranes were stained with an ATF4 antibody (Cell Signaling, #11815, 1:1000) and anti-vinculin antibody (Sigma Aldrich, #V9131) as a control. Secondary antibodies coupled to IR dye800CW, and IR dye680RD were employed for detection of HIF1-α, β-actin and vinculin. , whereas ATF4 was detected using horseradish peroxidase-coupled antibody. Membranes were imaged using the ChemiDoc™ MP Imaging System (BioRad). Quantification of band intensities was performed using Image lab Version 5.2 (BioRad).

**Quantification and statistical analysis**

All biological replicates were either representative of three independent (biological) replicates or expressed as mean ± SD. All statistical details of the conducted experiments can be found in the respective figure and table legends. n: number of biological replicates.

**Data availability**

The proteome datasets generated during this study are available at MassIVE (MSV000093287).
